# Experience-dependent development of dendritic arbors in mouse visual cortex

**DOI:** 10.1101/755934

**Authors:** Sarah E.V. Richards, Anna R. Moore, Alice Y. Nam, Shikhar Saxena, Suzanne Paradis, Stephen D. Van Hooser

## Abstract

Despite the importance of dendritic arbors for proper neuronal function, our knowledge of how sensory experience influences these structures during postnatal cortical development is incomplete. We present a large-scale dataset of 849 three-dimensional reconstructions of pyramidal neuron basal arbors collected across early postnatal development in the mouse visual cortex. We found that the basal arbor underwent a 45% increase in total length between postnatal day 7 (P7) and P30. Surprisingly, comparisons of dark-reared and typically-reared mice revealed that only 15% of arbor length could be attributed to visual experience. Furthermore, we characterized the role of the activity-regulated small GTPase Rem2, showing that Rem2 is an experience-dependent negative regulator of dendritic segment number during the visual critical period. These data establish a detailed, quantitative analysis of the basal arbor that has high utility for understanding circuit development and providing a framework for computationalists wishing to generate anatomically accurate neuronal models.

## Introduction

Dendritic architecture plays a fundamental role in the functional capacity of neurons [1–22]. Dendrites provide the real estate for housing synaptic contacts and, in concert with intrinsic membrane properties, dictate how incoming current will flow from these contacts to the soma. Despite the importance of dendritic architecture, much remains to be understood about how dendrites develop in mammalian sensory areas, the genetic programs that orchestrate this process, and how sensory experience may play a role. While many developmental studies have illuminated the details of cell migration [23–26], differentiation [23, 27–29], and dendritic spine formation [30–37], a high-resolution examination of the postnatal maturation of dendritic branches in a mammalian model system is lacking.

Existing reports suggest that the basal arbor of pyramidal neurons in visual cortex increases in length early in postnatal development and may reach a steady length beginning at postnatal day 30 (P30) [36, 37], at which time the arbor is largely resistant to changes in visual experience [38]. However, these works have focused on very early postnatal development [37] or sampled only a few later ages [36], which limits possible conclusions. More recent work has focused on post-critical period plasticity [39–41], leaving careful examination of dendrite development and plasticity in juvenile mice relatively unexplored. Interestingly, reports in other vertebrate species such as *Xenopus Laevis*, suggest that visual experience plays an important part in proper development of the dendritic arbor of optic tectum neurons and many molecular regulators of this process have already been identified [42–51]. However, numerous outstanding questions remain to be answered in mammalian systems: What are the developmental processes by which dendritic arbors adopt their adult configuration? Is maturation regulated by visual experience? Which molecular mechanisms underlie these processes?

To answer these questions, we performed a detailed analysis of dendritic architecture in layer 2/3 of mouse visual cortex at 5 ages beginning before eye opening (P7) and continuing through the height of the visual critical period (P30) [52]. Our analyses reveal that the length of the pyramidal cell basal arbor undergoes a significant outgrowth event between P16 and P21, and that this growth is composed almost entirely of increases in the length of dendritic segments but not an increase in the number of segments. That is, the net number of dendritic branches is relatively established by P7. Furthermore, we found that this increase in the length of dendritic segments is mediated by both experience-dependent and experience-independent processes. For the first time, we also establish Rem2 – a small GTPase previously implicated in dendritic complexity *in vitro* and in *Xenopus Laevis* optic tectum [51, 53–55] – as an experience-dependent negative regulator of dendritic complexity in mammalian visual cortex. This work demonstrates that Rem2 cell-autonomously regulates the arrangement of the basal dendrites: neurons in which Rem2 has been deleted are less likely to exhibit a significant directionality and sometimes exhibit abnormal directionality. Taken together, our results establish a unified framework for investigating dendritic development in the mammalian cortex and expand our understanding of how dendritic arbors are established and maintained in an experience-rich environment.

## Methods

### Golgi-Cox Labeling and Tissue Preparation

Typically-reared WT and *Rem2^-/-^* littermate mice were housed in a 12hr light/12hr dark cycle from birth until the specified age (P7, P12, P16, P21, P30). Dark-reared mice were placed in a light-tight box beginning at P9 until termination of the experiment (P16, P21, or P30). At the specified age, mice were anesthetized with ketamine/xylazine cocktail (ketamine 100mg/kg, xylazine 10mg/kg) and transcardially perfused with 0.9% saline in ddH20. Dark-reared mice were anesthetized in the dark and then shielded from light until after perfusion using a mask constructed of several layers of light blocking tape (Thor Labs) to cover the eyes. Immediately following perfusion, brains were weighed and submerged in Golgi-Cox solution (FD NeuroTechnologies). Throughout all steps involving Golgi-Cox, brains were protected from light. Golgi-Cox solution was changed 24 hours after initial immersion and brains continued to be stored in Golgi-Cox solution for 7 days. After 7 days, brains were transferred to Solution C (FD NeuroTechnologies) for at least 2 days. Coronal sections were cut at 150 µm using a cryostat and immediately mounted on slides coated with 2% porcine gelatin. Histology was carried out according to the protocol supplied by FD NeuroTechnologies RapidGolgi Stain Kit. Briefly, slides were washed with ddH2O, developed using FD Neurotechnologies Solutions D & E, rinsed in ddH2O, dehydrated with a graded series of ethanols, and cleared using xylenes. Slides were then coverslipped using Permount (Fisher Scientific).

### TdTomato Reporter Tissue Preparation

Typically reared *Rem2^+/+^;TdT^flex/flex^* and *Rem2^flx/flx^;TdT^flex/flex^* mice were housed in 12hr light/ 12hr dark cycle from birth until the specified ages and days post injection (dpi) (5dpi/P25, 7dpi/P27, 10dpi/P30). Mice (both *Rem2^+/+^; TdT^flex/flex^* and *Rem2^flx/flx^; TdT^flex/flex^*) were injected with AAV.GFP-Cre at P20 (see detailed method below) to activate reporter expression and, in *Rem2^flx/flx^; TdT^flex/flex^* mice, manipulate Rem2 expression. At the specified age, mice were anesthetized with ketamine/xylazine cocktail (ketamine 100mg/kg, xylazine 10mg/kg) and transcardially perfused with 0.1M Phosphate Buffered Saline (PBS) followed by 10% formalin in PBS. Immediately following perfusion, brains were extracted and transferred to 10% formalin for 24hrs at 4°C. Brains were then transferred to 30% sucrose in PBS for at least 2 days prior to sectioning in the coronal plane at 200 µm on a freezing microtome. Sections were immediately mounted on SuperFrost Plus (Fisher Scientific) slides, allowed to dry until adhered to slides, and covered with Fluoromount G (Diagnostic BioSystems, Inc.) and left to dry overnight. Slides were sealed with clear nail polish (Electron Microscopy Sciences). During all steps, the brains and brain sections were shielded from light whenever possible. Neurons were visualized using the genetically encoded TdTomato fluorophore and no additional staining of any kind was performed.

### Virus Injections

The AAV-GFP-Cre (AAV1.hSyn.HI.eGFP-Cre.WPRE.SV40; AV-1-PV1848) construct was obtained from the Vector Core facility at the University of Pennsylvania. Mice age P20 were anesthetized with a cocktail of ketamine (100 mg/kg) and xylazine (10 mg/kg), mounted on a stereotaxic frame, shaved, and the head was scrubbed using alternating washes of betadine and 70% ETOH. A single midline incision was performed, and underlying tissues scraped to expose the skull. Primary visual cortex was targeted by using the mouse brain atlas after adjusting for the lambda-bregma distance for age. In our hands, coordinates were approximately 3.5 mm lateral from the midline and 1.2 mm anterior to lambda. A small 1-2 mm in diameter hole was drilled in the skull and a glass micropipette delivered 50 nL AAV (diluted 1:250 in 0.9% sterile bacteriostatic saline) 150 μm below the dural surface. The incision was sutured, and triple antibiotic ointment was applied topically. Mice recovered on a heating pad and were returned to the animal facility until use.

### Neuron Selection and Imaging

Primary visual cortex was identified using landmarks, primarily the morphology of the hippocampus and corpus callosum, as depicted in the coronal Allen Reference Atlas (http://mouse.brain-map.org/static/atlas). Pyramidal neurons were selected from Layer 2/3 of primary visual cortex for reconstruction and analysis. Pyramidal neurons were identified by characteristic soma shape and the presence of a distinct apical dendrite oriented toward the cortical surface. Neurons selected for reconstruction met several criteria: the entire basal arbor was present in one tissue section, Golgi-staining or fluorescent fill was dense and complete in all processes, dendritic branches were not broken or truncated, artifact in the surrounding neuropil was minimal and allowed for unambiguous reconstruction.

A z-stack of images with a 2 µm step was captured of the entire basal arbor of each neuron using a 20X air objective. All Golgi-stained neurons were imaged using a Zeiss AxioObserver microscope under bright field illumination. Neurons expressing the TdTomato reporter construct were imaged using either a Nikon Eclipse Ti3 microscope or a Zeiss 880 LSM. For determining the medial-lateral orientation of TdTomato-expressing neurons, additional low-magnification images were captured with a 4X or 10X air objective using either a Zeiss 880 LSM or a Keyence BZX-700 microscope.

### Reconstruction of Neurons

All neurons were reconstructed using Knossos, a free annotation program developed by the Max Planck Institute for Medical Research (Heidelberg, Germany; https://knossos.app/). Images were prepared for reconstruction by unstacking each z-stack and converting to 8-bit tif files using a custom FIJI macro to be compatible with Knossos. Images were then converted to a Knossos-compatible format (called “cubing”) using Python code distributed by the creators of Knossos.

Neurons were reconstructed by manually placing sequential nodes in the center of the width of each dendritic branch along the entire basal arbor. Neurons were reconstructed in 3 dimensions by scanning through the depth of the z-stack and placing nodes where each segment of dendrite was in focus. Care was taken to conform to the curvature of dendritic branches by increasing node density in highly tortuous regions. All dendritic primary arbors were connected to a central, spherical node at the soma that was sized to match the diameter at the widest point of the soma (used as calculation of soma size as approximated as a sphere).

### Analysis of Reconstructions

Neuronal morphology was analyzed using a suite of custom Python code created for this project. The analysis code relies on identities and 3D coordinates of each node and associated auto-generated edges to calculate a variety of parameters about whole dendritic arbors as well as individual primary arbors. Detailed explanations of each analysis are found in Table 1. All code is available on the Van Hooser Lab GitHub repository (http://github.com/VH-Lab/) in the RichardsEtAl_Original project folder. A more-streamlined version of this code, ideal for adaptation to similar projects for morphology quantification, is also provided (RichardsEtAl_Streamlined). See more detailed descriptions for complex measures below. Additional measures not used in this publication are described in detail in the README included in the Python project for these analyses.

**Table 1:**
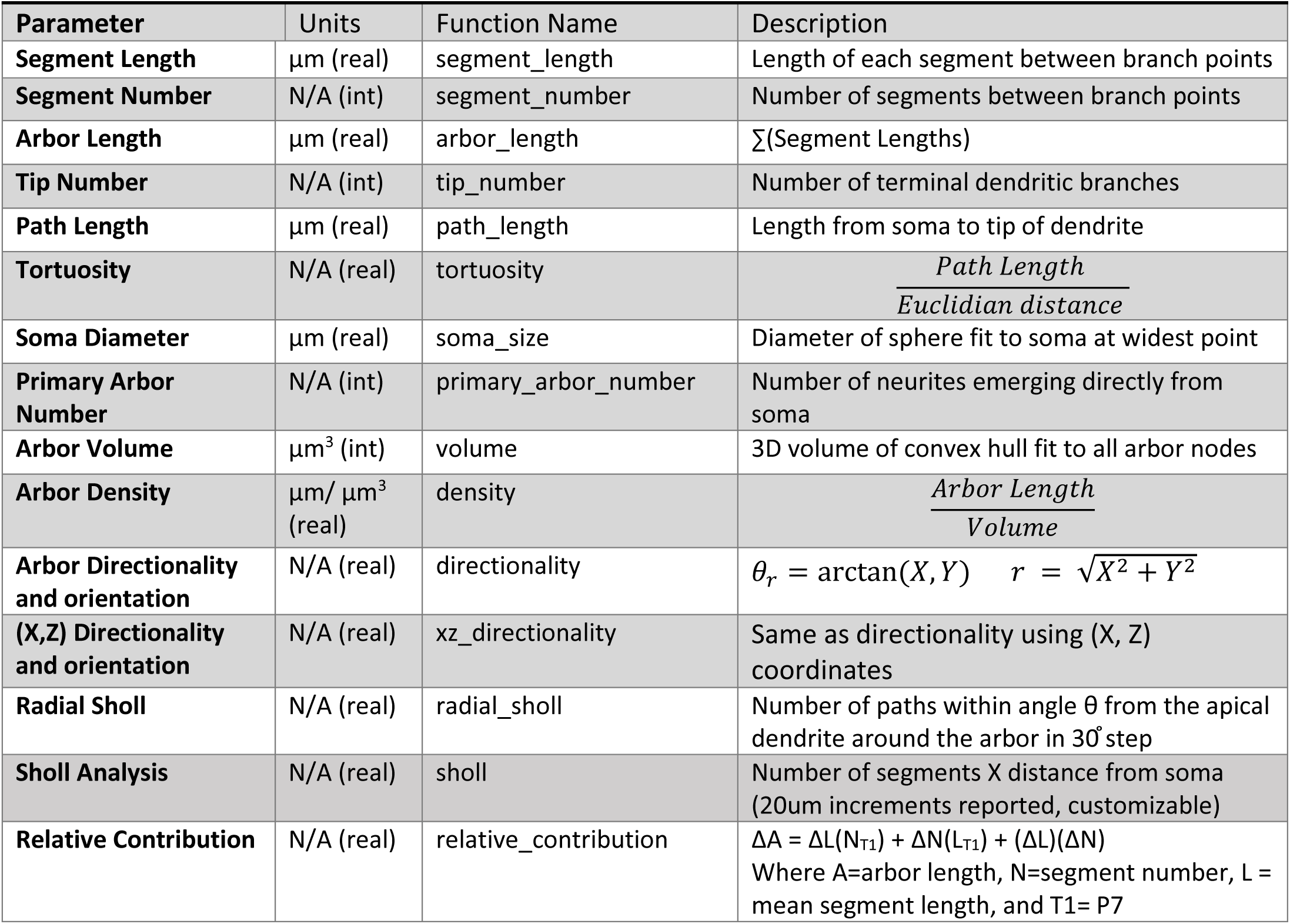
Analyzed Parameters. List of each parameter name, unit of measure, function name in custom analysis package, and plain language description of measurement.

Arbor length changes over time (ΔA) due to changes in the number of branches (ΔN), changes in the length of existing branches (ΔL) and the combined contribution of these factors see Figures 3E and 6E). We computed the fractional contribution of each of these components between each pair of neurons in each group (e.g. all P7 neurons compared to all P12 neurons, and so on) as below, where N_1_ and L_1_ indicate the initial time point and N_2_ and L_2_ indicate a subsequent time point (e.g. to measure developmental change from P7 to P12, P7 is the initial state and P12 is state 2)

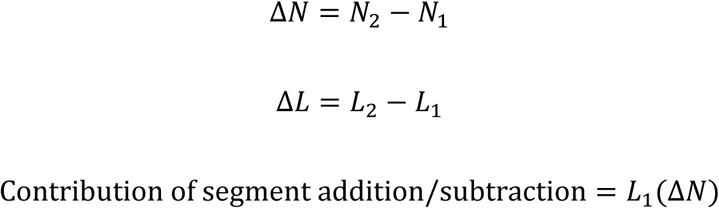

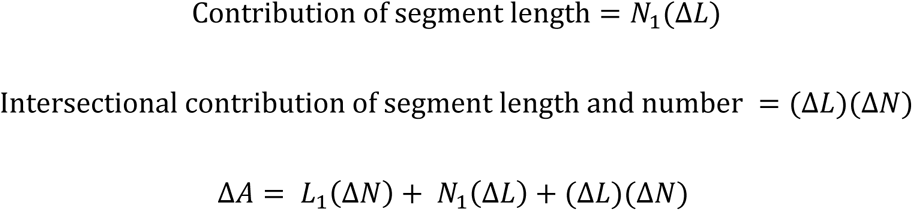

A value of 0 for any measure indicates no change in arbor length results from that measure. Positive values indicate that changes in a given measure contributed to increased arbor length were as negative values indicate that changes in a given measure lead to decreased total arbor length.

To calculate basal arbor orientation and directionality, neuronal reconstructions were rotated to align the axis of the apical dendrite to 0°. Because neurons were reconstructed from both hemispheres, some reconstructions were reflected across the Y axis so that the lateral side of each arbor was at 90° and the medial side at 270°. The tip node of each path was used to create vector between the path tip and the soma node with length of 1 unit and the angle (*a*) between each path vector (*path*) and the apical vector (*apical*) was used for calculation of the mean vector of path angles. The degree of orientation or directionality is indicated by the length of the mean vector (r), which ranges from 0-1. The vectorized arbors were also used to calculate Radial Sholl and Apical Cone Measures (see Table 1)

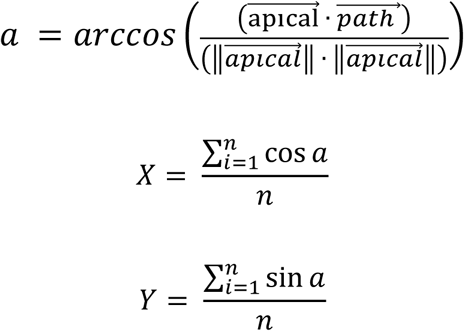

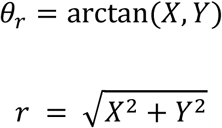

Significant orientation and directionality was determined by calculating the Raleigh Z statistic and comparing this value to the critical value for significant directionality for each arbor where *n* is the number of paths and *r* is the mean vector.

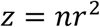

## Results

### Quantification of Arbors

In order to quantify the development of basal dendritic arbors in mouse visual cortex, we developed a reconstruction and analysis pipeline consisting of freely available reconstruction software (Knossos, Max Planck Institute, Heidelberg) and a library of custom analyses written in Python. Figure 1 describes the majority of the analyses as performed on an example cell (Fig. 1A). Our analyses can best be grouped into arbor-centric (Fig. 1B-E, H) and tissue-centric (Fig. 1F, G, I) measures. Arbor-centric measures focus on characterizing each arbor in terms of its size (Fig 1C), number of components (Fig. 1D-E), and measurements of individual components (Fig. 1 B&H). In contrast, tissue-centric measures characterize how the arbor is arranged with respect to the entire tissue, such as the volume the arbor occupies in the cortex (Fig. 1G) or how the dendrites are arranged in the space around the soma (Fig. 1F). Both types of measures consider the arbor as three-dimensional when relevant, such as when calculating length-based parameters.

**Figure 1:**
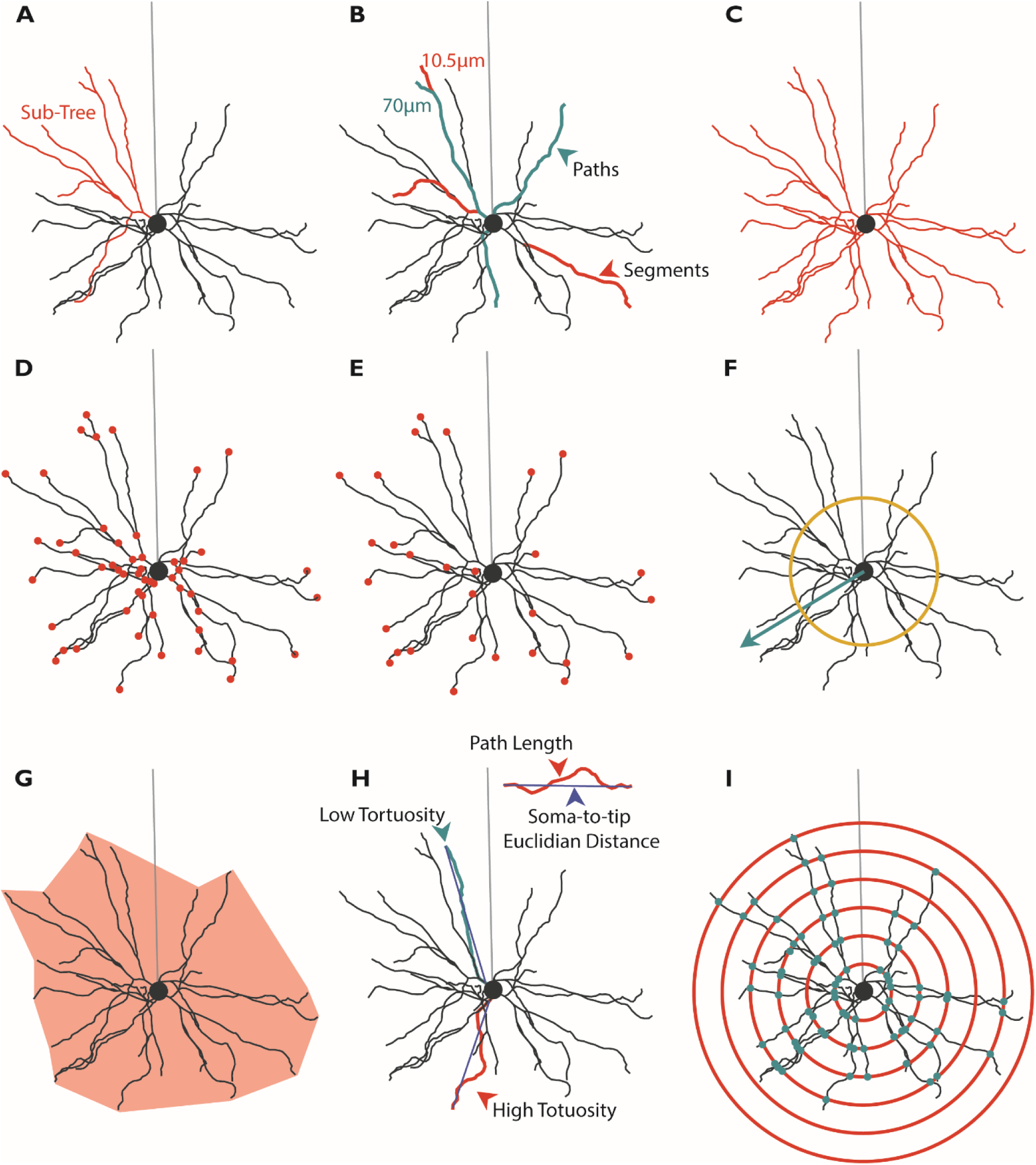
Analysis of basal dendrite morphology. **A)** Example hand-reconstructed L2/3 wildtype pyramidal neuron sampled from mouse primary visual cortex. One primary arbor is highlighted in coral. Position of un-reconstructed apical dendrite indicated in light gray. Soma is diagramed as a circle. **B)** Segment (coral) and Path (teal) diagram. Segments are the smallest span of dendrite between two branch points. Paths are the length of dendrite that connects branch tips to somas. Paths are composed of one or more segments. **C)** Total Arbor Length diagram. Total arbor length is the summed length of all segments, measuring the total length of the arbor in microns. **D)** Segment Number diagram. Each dot (coral) indicates one branch point in the arbor. **E)** Tip Number diagram. Each dot (coral) indicates the terminal tip of a path. This is equivalent to the number of paths. **F)** Arbor Polarity diagram. Arbor polarity is calculated in several way including analysis of dendritic material (number and length of segments and paths) on each side of the soma and in terms of absolute arbor directionality calculated as mean vector length (teal arrow) and tested using the Raleigh Test. Mean vectors with length greater than yellow ring indicate significant directionality of the arbor. **G)** Arbor Volume diagram. Arbor volume is the 3-dimensional volume of a least-size polygon fitted to all points in the basal arbor (light coral). Arbor density is the Total Arbor Length divided by the Arbor Volume. **H)** Tortuosity diagram. Tortuosity is the Path Length for an individual path divided by the soma-to-tip Euclidian distance. **I)** Sholl analysis diagram. Sholl analysis counts the number of segments (teal dots) at several evenly-spaced distances from the soma (coral circles) to measure the relationship between distance from the soma and arbor complexity.

Measures also differ in whether the unit of dendritic length is the dendritic segment or the dendritic path. Dendritic segments (Fig. 1B) are the smallest units of dendritic material and are defined as the length of dendrite spanning two branch points, between the edge of the soma and a first branch point, or between a branch point and a tip. Dendritic paths (Fig. 1B), in contrast, are the summed length of dendrite between the edge of the soma and the branch tip (Fig. 1E). A dendritic path can be conceptualized as the distance a synaptic current originating at the branch tip (Fig. 1E) would have to travel to reach the soma. Given this definition, dendritic paths are typically an assembly of several dendritic segments, except in the case of an unbranched segment which is considered to be a dendritic segment, path, and primary arbor. Primary arbors (Fig. 1A) are the largest sub-assembly of the arbor and are defined as a primary neurite emerging from the soma and all subsequent dendritic segments arising from that primary neurite.

### Setup and Initial Characterization

Previous studies have quantified dendritic arbor development in mouse primary visual cortex using very sparse sampling during development or have analyzed dendritic arbors at times that are well past the relevant epochs of circuit plasticity such as the extensively-characterized critical period for ocular dominance plasticity (Fig. 2A, top)[36, 37]. We set out to densely sample arbor development (Fig 2A, bottom) by assessing arbor complexity approximately every five postnatal days starting before eye opening (postnatal day 7 (P7)), at eye opening (postnatal day 12 (P12)), and during the first few days of visual experience (postnatal day 16 (P16)). We also made measurements near the opening of the critical period (postnatal day 21 (P21)) and during the peak of the critical period (postnatal day 30 (P30)). To visualize neurons, we performed Golgi-Cox histology and identified L2/3 pyramidal neurons based on canonical morphological features of pyramidal neurons, including a triangular cell body and apical dendrite oriented toward the pial surface. All reconstructed neurons had complete basal dendritic arbors and were located in primary visual cortex (Fig 2B). We then made manual reconstructions using Knossos (Max Planck Institute for Medical Research; Heidelberg, Germany) and analyzed them as described above (Fig. 1 and Table 1).

**Figure 2:**
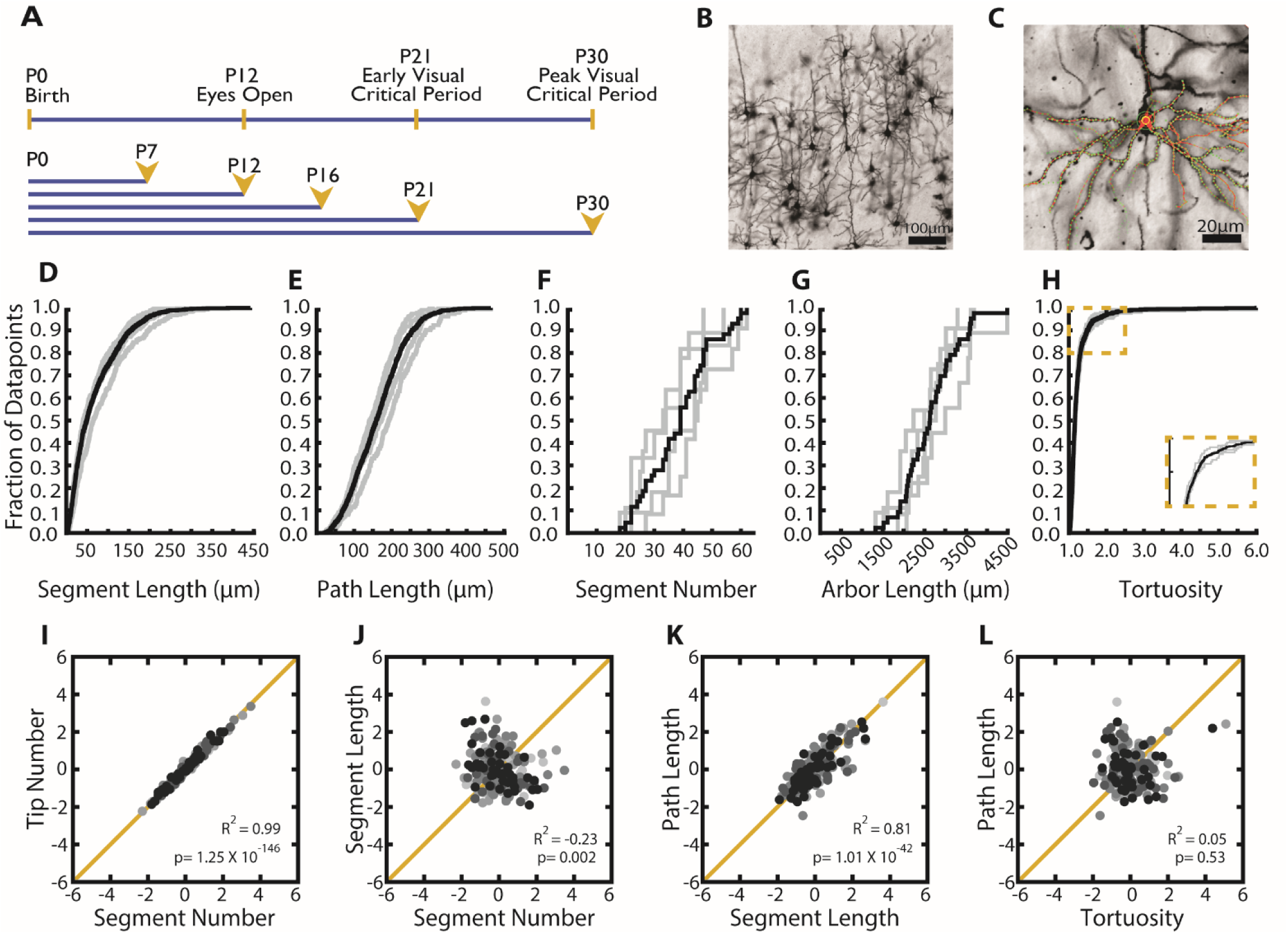
Distributions of observed values for arbor parameters suggest distinct regulatory mechanisms. **A)** Timelines of visual development in the mouse (top) and morphological sampling across development (bottom). Ages sampled indicated by postnatal day (P#) and yellow arrowheads. **B)** Example field of Golgi-stained cortical pyramidal neurons demonstrating dark fill of neurites and ability to differentiate cell types. **C)** Example 3D tracing of basal dendrites of L2/3 pyramidal neuron in visual cortex. Green circles indicate experimenter-placed nodes. Red lines are edges connecting nodes. Some traced branches are not in-plane. **D-H)** Cumulative distributions of **(D)** Segment Length **(E)** Path Length **(F)** Segment number **(G)** Arbor Length and **(H)** Tortuosity. Gray lines are all observations from individual wildtype mice at age P30. Black lines are the complete population of all observations across animals at P30. **I-L)** Correlations between listed parameters for entire population of all neurons across all ages. R^2^ values and p-values are listed for each comparison. Neurons sampled at different ages are indicated by different colors with light greys indicating young animals and black indicating older animals. No noticeable clustering by age was observed.

**Figure 3:**
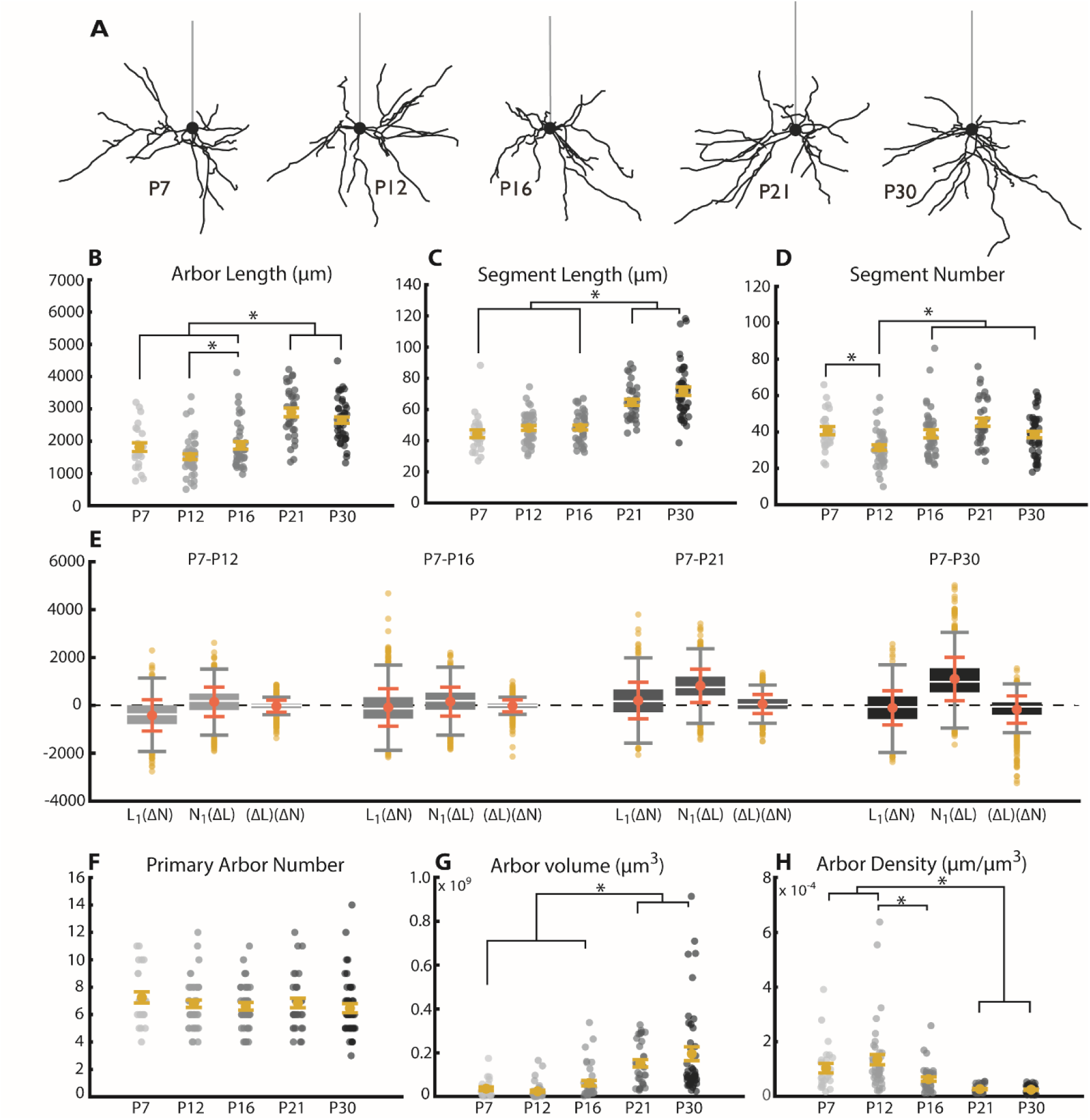
Length and volume of wildtype arbors undergo significant growth near the opening of the visual critical period coincident with an increase in the length of dendritic branches. **A)** Example reconstructions of basal arbors from L2/3 wildtype visual cortex neurons at the five ages sampled. Straight gray lines diagram the unreconstructed apical dendrite to indicate the orientation of cells. **B)** Total arbor length of the basal arbor across development (light to dark grey dots are individual neurons). **C)** Segment length of basal dendrites across development. Segment length for each neuron is an average of all path lengths sampled from that cell (light to dark grey dots are individual neurons). **D)** Segment number of basal dendrites across development (light to dark grey dots are individual neurons). **E)** Analysis of relative contribution of changes in segment number (L_1_(ΔN)), segment length (N_1_(ΔL)), or the combination of these factors ((ΔL)(ΔN)). Boxes are 25-75^th^ percentile of the data. Median is indicated by a white line. Mean and standard error are indicated by coral dot and error bars. **F)** Number of dendritic trees (number of primary neurites) of basal dendrites across development. Flyers are indicated by yellow dots. **G)** Three-dimensional volume of convex hull fit to nodes composing basal arbor across development. **H)** Density of the basal arbor across development. Calculated as microns of dendritic material per cubic micron of arbor volume. Error bars are mean ± SEM. Yellow dots indicate the mean and gray dots indicate single neurons. For all measurements: P7 N=24 cells, 2 mice; P12 N=43 cells, 4 mice; P16 N=38 cells, 4 mice; P21 N=33 cells, 3 mice; P30 N=43 cells, 4 mice (* indicates P<0.05, 1-way ANOVA with post-hoc Tukey).

In order to establish organizing principles of dendritic arbors, we performed analyses on the entire population of reconstructed neurons at a given age, in this case P30, both between animals (individuals animals are gray lines) and pooled across animals (black Lines). We found that observed values of each measure show a unique amount of variability in terms of both total variation (Fig. 2D-H, black lines) and between-animal variation (Fig. 2D-H, grey lines). For example, we found a wide range of dendritic segment counts across the population as well as between animals (Fig. 2F). Conversely, we found that the length of these dendritic segments and paths is relatively less variable both on the whole and across animals (Fig. 2D-E). This suggests that even though the number of paths is quite variable, their length is fairly constant, which may suggest distinct regulatory mechanisms of these two characteristics. As might be expected, the length of whole arbors (Fig. 2G, sum of length of segments) is intermediately variable compared to the number and length of paths (Fig. 2 D-F). Interestingly, some measures of arbor complexity, such as tortuosity (Fig. 1H and Fig. 2H), are extremely invariant both within and between animals. This may indicate strict maintenance of a set value either through precise regulation at the molecular level or unknown biophysical constraints that tightly restrict path tortuosity.

We observe that several neuron-centric parameters are highly correlated. As expected, the number of branch tips is almost directly proportional to the number of dendritic segments (Fig. 2I), which led us to focus only on segment number for the remainder of this article. Similarly, the length of dendritic paths is very closely related to the length of segments (Fig. 2K). While these relationships may be expected, we also found novel correlations between measures. For example, we found that segment number is negatively correlated with segment length (Fig. 2J), suggesting that arbor length overall may have a set point and the architecture of the arbor develops to fit this loosely pre-determined total length. Interestingly, we at times found no relationship between quantities that on first consideration may seem likely to be co-regulated, such as the length of paths and their tortuosity (Fig. 2L). Taken together, these results suggest that some arbor parameters may be co-regulated, while others may be regulated by isolated processes.

### Development of Wildtype Arbors in Typical Rearing Conditions

In wildtype cells, the total length of the basal arbor tends to increase over time with some additional nuanced dynamics (Fig. 3A). Early in development, between P7 and P16, arbor length is relatively stable with the exception of a small-but-significant increase in arbor length between P12 and P16 (Fig. 3B). However, there is a large increase in total arbor length between P16 and P21, resulting in P21 arbors that are significantly larger than arbors at any of the three earlier time points (Fig. 3B). When we observed arbors at P30, we saw no change from arbors at P21 (Fig. 3B), suggesting that total arbor length may have stabilized. This is supported by previous work indicating that arbors are stable from P30-P60 in rodent visual cortex [36, 37].

When considering how this increase in total arbor length could arise, three possible sources of new arbor are immediately evident – existing segments increase in length, new segments are added, or a combination of these two factors accounts for the added length. To differentiate among these possibilities, we quantified each factor. Similar to the total length of the arbor, we found that the mean length of individual segments undergoes a sharp increase between P16 and P21 resulting in an increase in the average segment lengths at P21 compared to all earlier time points (Fig. 3C). No further increase is seen between P21 and P30, as was also observed with total arbor length (Fig. 3C).

However, a different pattern of developmental change emerged when we instead considered the number of segments that compose the arbor. During pre-critical period development, we observed small, dynamic changes in segment number characterized by an early reduction in mean segment number between P7-P12 followed by an increase in mean segment number from P12-P16 (Fig. 3D). Segment number during the critical period (P21-P30) is similar to that observed at P7 or P16 and differs significantly only from P12 (Fig. 3D), highlighting the lack of a distinct transition point that we observed in both total arbor length and segment length. While there was a noticeable numerical decrease in mean segment number between P21 and P30, this difference did not reach the criterion for statistical significance (p=0.09). Interestingly, the number of primary arbors did not change during postnatal development (Fig. 3F), which suggests that arbor-centric development occurs primarily through regulation of segments as opposed to the addition or removal of primary arbors.

To better understand whether the outgrowth of existing segments (Fig. 3C) or the addition of new segments (Fig. 3D) contribute more to arbor development, we computed the relative contribution of these factors (Fig. 3E), as well as their interaction, to the observed changes in total arbor length (Fig. 3B) between each pair of neurons at each age compared to our youngest age, P7 (Table 1). We found that changes in segment number impact the small change in arbor length most in the P7-P12 comparison, but that changes in segment length predominated at each subsequent age (Fig. 3E). This effect is most clearly demonstrated at the later time points we investigated (P7-P21 and P7-P30), at which point changes in segment number account for effectively no change in arbor length (Fig. 3E). Our results strongly suggest that overall arbor length arises predominantly through changes in the length of dendritic segments, with changes in segment number being a minor component primarily influencing early development.

In addition to developing characteristics within the arbor of a single cell, it is worth considering whether the “footprint” of the arbor within the larger tissue changes over time. To quantify this, we investigated both the volume that the arbor occupies within the tissue as well as the density of the arbor within that area. The volume, or 3D spread, of the arbor relates to the area over which the arbor can search for synaptic partners whereas the density is related to the likelihood of a synapse occurring at any given point in that space.

As arbors grow and change, the “footprint” they occupy in the larger tissue may also be altered, impacting the availability of synaptic partners and the likelihood that a given axon will encounter the arbor if passing through a region of cortex. When we considered arbor volume (Fig. 3G), we observed that this feature corresponds to arbor length extremely well. Arbor volume undergoes a sharp transition between P16 and P21 that is maintained through P30 (Fig. 3G). Also notable is the remarkable increase in variability over development. In contrast, arbor density decreased over development (Fig. 3H), with the most striking decrease occurring at P21, similar to the timing of arbor volume development. This indicates that, even though both arbor length and arbor volume are increasing, the three-dimensional spread of the arbor outpaced the increase in arbor length, leading to decreased arbor density. Interestingly, the variability of arbor density starkly decreased throughout development, complementary to the variability increase seen in arbor volume (Fig. 3G-H). This indicates that mature arbors vary greatly in volume, perhaps indicating cell type-specific arbor volumes, but maintain very similar density across cells.

### Basal Arbors show Distinct Coronal and Tangential Orientation and Directionality

The relative placement of dendritic paths in space dictates the ability of presynaptic partners to encounter postsynaptic cells. To this end, tiling of sensory tissues by dendritic processes is a fundamental organizing principle of sensory circuits, such as in the mammalian retina. Several known molecular regulators, such as DSCAMs [56–59] and Semaphorins [60], contribute to the establishment of proper arbor shape, self-avoidance, and mosaic spacing in the mammalian retina. Though tiling has been hypothesized to be present in higher order sensory areas such as the visual cortex, a direct measurement of dendritic orientation or directionality has not been made in rodent systems and few molecular regulators have been identified [59]. Thus, we became interested in calculating basal arbor orientation and directionality and turned to circular statistics [61] commonly used to assess direction selectivity in visually-responsive neurons. In short, cells were aligned along the axis of their apical dendrite and a mean vector angle and length was computed from the angular orientation of each normalized path vector. Normalized vector length is expressed as a value ranging from 0 to 1 in which 0 indicates no directionality (or orientation) and 1 would indicate a perfectly directional (or oriented) arbor. We calculated basal arbor directionality and orientation in two viewpoints: a coronal viewpoint (Fig. 4A-I) as would be achieved using a standard coronal sectioning technique and a tangential viewpoint (Supplemental Fig.S1) as would be achieved by looking down onto the cortical surface. We computed directionality in neurons sampled from wildtype mice between 23 and 30 days of age (P23 – P30).

**Figure 4:**
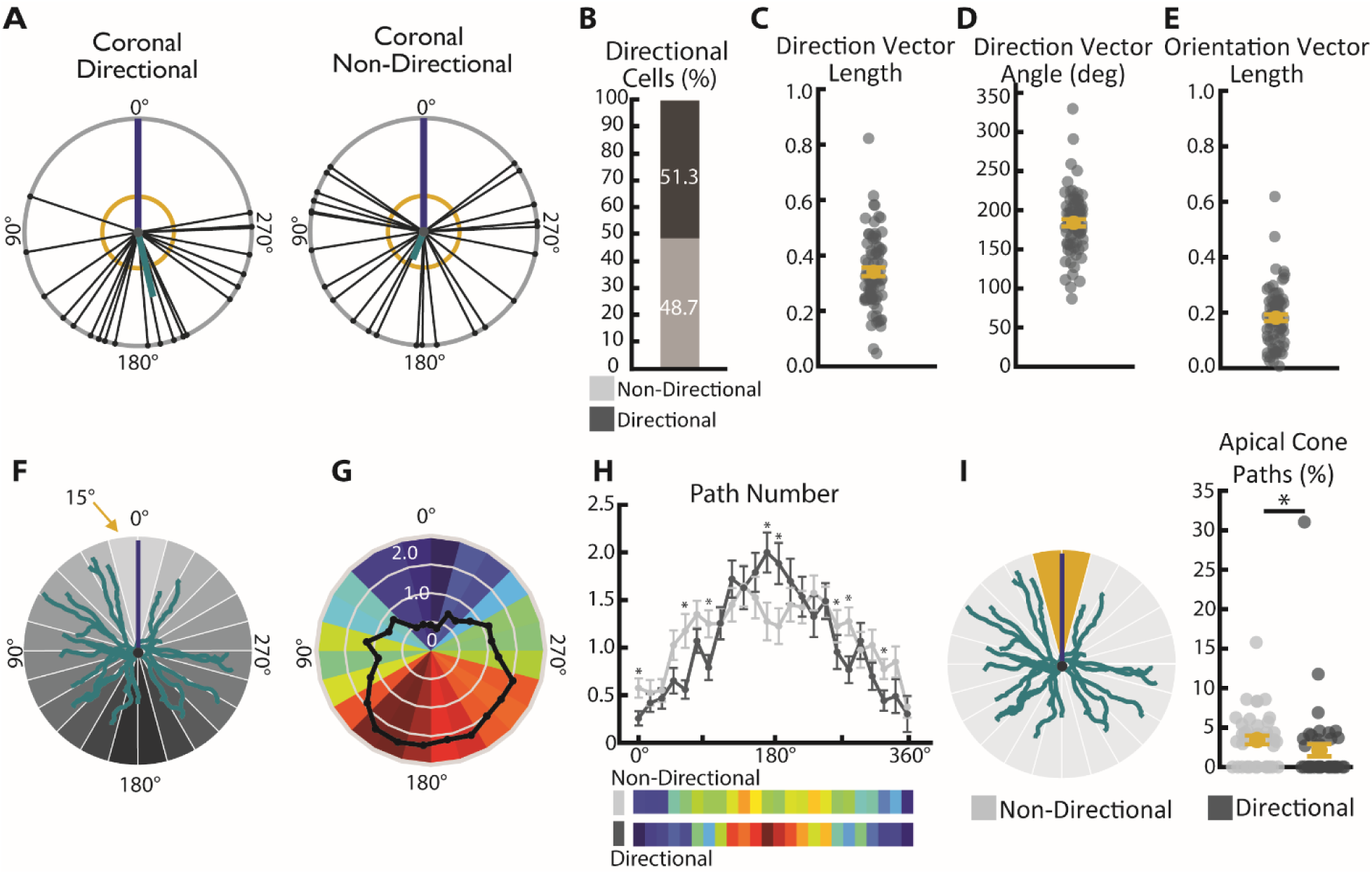
Basal arbors show significant directionality in path location. **A)** Example directional (left) and non-directional (right) basal arbors sampled from wildtype mice in the coronal viewpoint. Each arbor is schematized as soma (center dot) and soma-to-tip vectors (lines with dots at tip) arranged on a unit circle with apical dendrite diagramed as blue line. Thick teal line indicates the mean directional vector and yellow circle indicates length of mean directional vector required to declare the arbor directional by the Raleigh Test. **B)** Pie chart showing portion of total wildtype population that are categorized as direction and non-directional using the Raleigh Test. **C)** Length of the coronal normalized directional vector calculated from all basal paths. Larger values indicate greater directionality. **D)** Angle of coronal directional vector calculated from all basal paths. **E)** Length of the coronal normalized orientation vector calculated from all basal paths. Larger values indicated greater orientation. **F)** Diagram of Radial Sholl measure. Number of paths falling into each 15 degree increment (grey wedges) is calculated for each cell. Basal arbor diagrammed in teal, apical dendrite position diagrammed in blue. **G)** Heat map indicating number of paths in each 15 degree bin for each cell (colorful wedges) from many (warm colors) to few (cool colors) and the mean (black line) **H)** Diagram of apical cone measurement. The number of paths falling in the 30 degree span around the apical dendrite (“apical cone,” yellow wedges) is calculated for each cell. **I)** Number of paths in the apical cone in significantly directional (dark grey) and non-directional (light grey) arbors.

In the coronal plane, we found that slightly greater than 50% of neurons have significantly directional arbors as determined by Raleigh Z Test. The directional vector of these arbors tends to be at 180°, or directly opposite the apical dendrite (Fig. 4D). In order to gain a more nuanced view of path location, we performed additional analyses in which the 360° around the soma were divided into 15° bins and the number of paths falling into each bin was calculated (Fig. 4F) for each neuron. When the number of paths was averaged across the population, we observe an enrichment of paths near 180° and a paucity of paths falling near the apical dendrite (Fig. 4G). When we compared directional and non-directional cells as determined by the Raleigh Z Test, we found a widening of the distribution of paths such that non-directional cells have an increase in the number of paths falling in the approximately 0-90° and 270-360° regions as well as a decrease in the number of paths within the region adjacent to 180° (Fig. 4H). We also found that non-directional cells show a small-but-significant increase in the number of paths falling within the region directly adjacent to the apical dendrite, which we have termed the “apical cone” (Fig. 4I). Despite these small differences, basal arbors of all cells showed a strong tendency to avoid the “apical cone”. When we considered tangential directionality and orientation, we observed that that only 10-20% of arbors exhibit directionality (Fig. S1D), but that 100% of the arbors sampled demonstrated robust orientation along the medial lateral axis (Fig. S1B-C). However, we are unable to distinguish whether this principle applies to all L2/3 pyramidal neurons or only our target population, which consists entirely of cells whose whole basal arbors fit into relatively thin (150µm) histological sections. Further studies will be required to establish how this finding may generalize to larger L2/3 neurons that may have distinct arbor shapes from those observed here.

### Development of Wildtype Arbors in the Absence of Visual Experience

We were interested in understanding how visual experience might impact dendritic development in the mammalian visual cortex, because previous work has found visual experience to be an important regulator of dendritic maturation in other species [46, 48–50, 62–66]. To address this question, we used dark rearing to eliminate visual experience and sampled complexity at three time points corresponding to early visual experience (P16) the opening of the critical period (P21) and the peak of the visual critical period (P30) (Fig. 5A), at which point we have evidence that the arbor size and architecture is relatively stable (Fig. 3B). We observed no change in total arbor length between cells from typically-reared and dark-reared mice at the earliest time point (P16), but we did observe that arbors from dark-reared mice were smaller than those from typically-reared mice at both P21 and P30 (Fig. 5C). Interestingly, we found that arbors sampled from dark-reared mice have a greater total length at P21 and P30 than arbors from P16 mice (both typically-reared and dark reared, compare Fig.5C), indicating that some arbor growth occurs even in the absence of visual experience. This suggests that arbor growth in typically-developing animals is likely mediated by both experience-dependent and experience-independent mechanisms (Fig. 5C), possibly including activity-dependent negative regulators to limit outgrowth when visual experience is present [51]. However, our findings establish that the majority of postnatal arbor branching occurs independently of visual experience in the mouse visual cortex, with only about 15% additional arbor length added with experience, which is in sharp contrast to previous findings that sensory input strongly promotes both dendrite branching and length in *X.Laevis* tadpoles [46].

**Figure 5:**
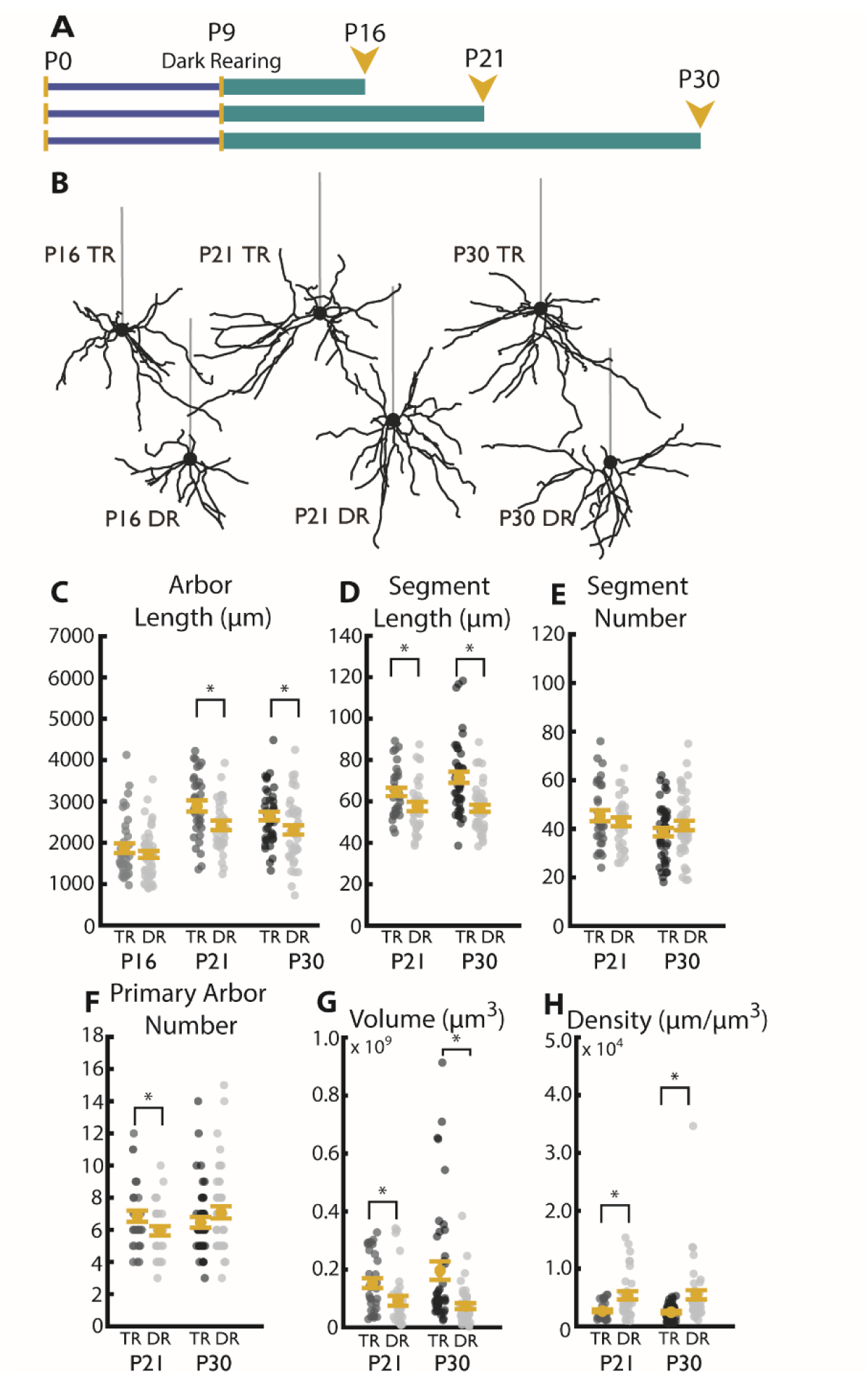
Visual experience during the critical period promotes increases in segment length leading to typical arbor development. **A)** Timeline of dark rearing experiments. All mice were dark reared starting at P9 and dark rearing (indicated by teal bar) continued until sacrifice at P16, P21, or P30. **B)** Example reconstructions of typically reared (TR, 12hr light/ 12hr dark) and dark reared (DR, 24hr dark) neurons at P16, P21, and P30. Straight gray lines diagram the unreconstructed apical dendrite to indicate the orientation of cells. **C)** Total arbor length with (TR) and without (DR) visual experience at P16, P21, and P30. **D)** Segment length at P21 in neurons sampled from mice with and without visual experience at ages P21 and P30. **E)** Segment number at P21 and P30 in neurons sampled from mice with and without visual experience. **F)** Primary arbor number sampled at P21 and P30 with and without visual experience. **G)** Arbor Volume sampled at P21 and P30 with and without visual experience. **H)** Arbor density sampled at P21 and P30 with and without visual experience. Error bars are mean ± SEM. Yellow dots indicate means and gray dots indicate single neurons. For all measurements: P16 TR N=38 cells, 4 mice; P16 DR N=46 cells, 4 mice; P21 TR N=33 cells, 3 mice; P21 DR N=32 cells, 4 mice; P30 TR N=43 cells, 4 mice; P30 DR N=45 cells, 4 mice (* indicates P<0.05, Wilcoxon rank-sum test).

**Figure 6:**
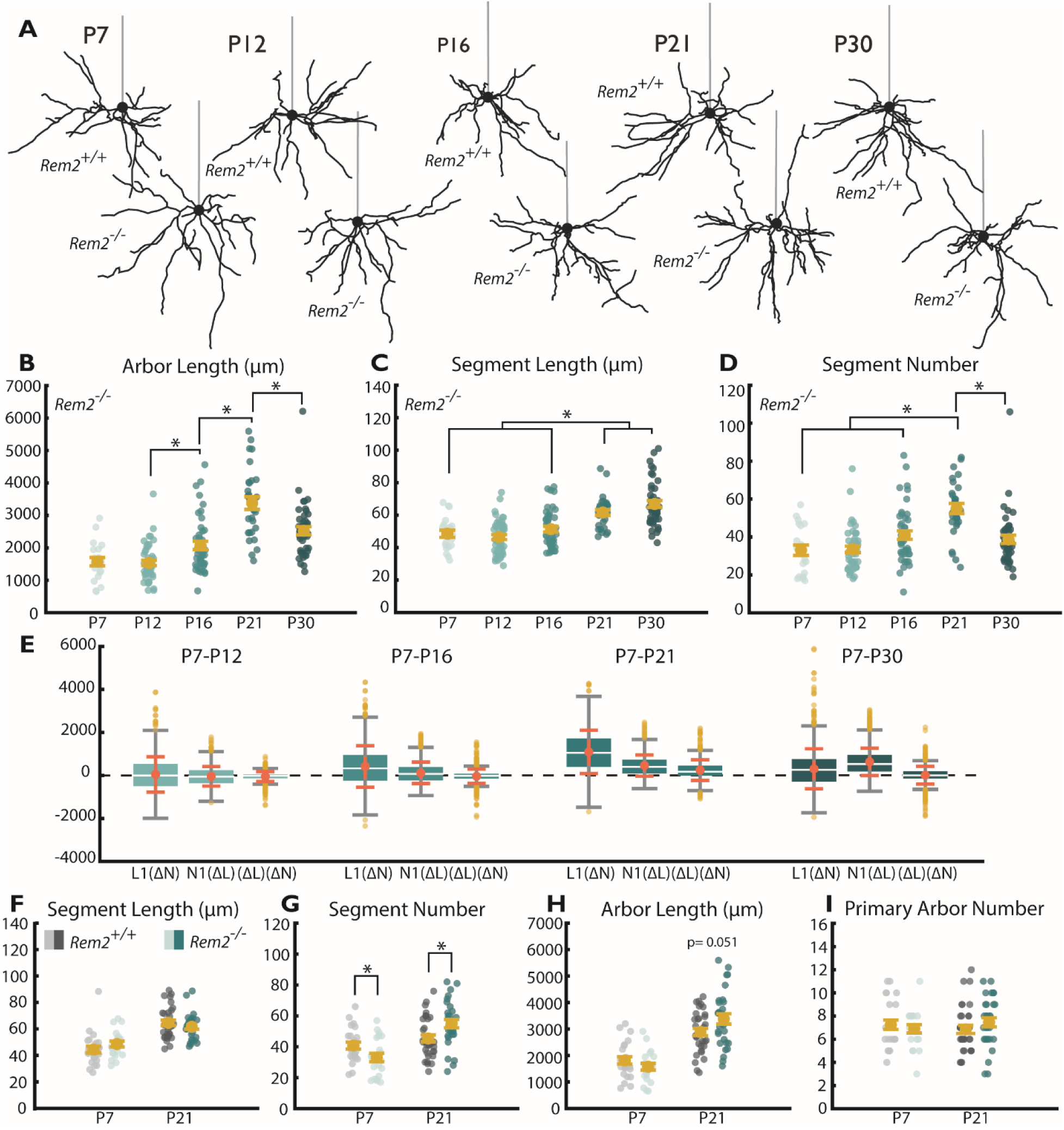
Rem2 regulates the development of basal dendritic arbors. **A)** Example reconstructions of typically reared (TR, 12hr light/ 12hr dark) wildtype (*Rem2 ^+/+^*) and Rem2 knockout (*Rem2^-/-^*) neurons at ages P7, P12, P16, P21, and P30. Straight gray lines diagram the unreconstructed apical dendrite to indicate the orientation of cells. **B)** Total arbor length of the basal arbor in *Rem2^-/-^* (teals) neurons at P7, P12, P16, P21, and P30. **C)** Average length of dendritic segments of *Rem2^-/-^*neurons sampled at P7, P12, P16, P21, and P30. **D)** Segment number of *Rem2^-/-^*neurons sampled at P7, P12, P16, P21, and P30. **E)** Number of dendritic primary arbors of basal dendrites across development. Flyers are indicated by yellow dots. **F)** Average segment length at ages P7 and P21 sampled from *Rem2 ^+/+^* (grays) and *Rem2^-/-^* (teals) neurons. **G)** Segment number at ages P7 and P21 sampled from *Rem2 ^+/+^*(grays) and *Rem2^-/-^* (teals) neurons. **H)** Total arbor length at ages P7 and P21 from *Rem2 ^+/+^* (grays) and *Rem2^-/-^*(teals) neurons. **I)** Primary Arbor number at ages P7 and P21 from *Rem2 ^+/+^* (grays) and *Rem2^-/-^* (teals) neurons.

The differences in dendritic arbor lengths with and without experience could arise from three general sources – a decrease in the number of segments, a decrease in the length of segments, or a combination of both. We observed no change in the number of segments (Fig. 5E) but found that arbors sampled from typically-reared animals exhibited significantly longer dendritic segments at both P21 and P30 (Fig. 5D) compared to dark-reared animals, indicating that visual experience is required to properly establish the length, but not number, of dendritic segments in wildtype mice.

Interestingly, we also observed marked changes in the “footprint” of each cell in the tissue. As expected given a decrease in arbor length, cells from dark-reared mice exhibited a decrease in arbor volume relative to those from animals with typical visual experience (Fig. 5G). We observe a complementary increase in density (Fig. 5H) in dark-reared cells, indicating that arbor length and volume do not necessarily scale to maintain arbor density. One possible outcome is that this basal architecture could decrease the variety of synaptic partners that a cell is able to access but maximize the likelihood of a partner axon in the region contacting a dendrite probabilistically. Stated differently, in the case the cortex receives visual input, arbors expand in both length and volume but decrease in density. While little is known about the possible functional impact of these changes, one might hypothesize that such an architecture may give greater access to diverse synaptic partners but with a lower chance of stochastically encountering any given partner.

### Rem2 Contributes to Arbor Development through Regulation of Segment Number

The findings presented here suggest that proper development of dendritic arbors progresses through interaction of experience-dependent and experience-independent mechanisms. Furthermore, our findings show that parameters of arbor complexity do not necessarily follow a single pattern of change or even demonstrate correlated changes, but instead may be regulated independently such that some values increase while others decrease or undergo no significant alteration over time at all. For example, when basal arbors grow from P7 to P30, they do not haphazardly increase length and add segments, but appear instead to specifically increase segment length as opposed to forming new branches. While observations such as these do not amount to direct evidence that segment number and length are governed by distinct regulatory programs at the molecular level, it does seem likely that these decoupled parameters would require some distinct regulatory mechanisms to orchestrate such discrete patterns of development. We were, therefore, interested to investigate whether a known regulator of dendritic complexity influences one of these defined characteristics of dendritic architecture (e.g. arbor length, segment number, tortuosity, etc.). We chose to investigate the small-Ras like GTPase Rem2, a known activity-dependent negative regulator of dendritic complexity identified both *in vitro* in dissociated cortical culture and *in vivo* in the *X. Laevis* optic tectum [51, 53–55]. To this end, we used Rem2 knockout littermates of our previously discussed wildtype mice to investigate the role of Rem2 in regulating the development of dendritic complexity.

To better understand how loss of Rem2 might impact the dendritic arbor of L2/3 pyramidal neurons in visual cortex, we assessed how development of the arbor progresses over time with regard to each of our key measures of dendritic complexity (Fig. 6A-D), finding changes in the trajectory of arbor length addition (Fig. 6B) and the number of dendritic segments (Fig 6D). We also calculated the relative contribution of segment addition, segment elongation, and the combination of these changes. In contrast to wildtype neurons (Fig. 3E) in which elongation of segments is the primary contributor to arbor length increases, segment number plays a more central role in the arbor development of *Rem2^-/-^* neurons (Fig. 6E, Fig. 6F-H). This is especially noticeable at the later time points, P21 and P30, which lacked the predominance of length-based changes characteristic of wildtype arbors (compare Fig. 3E to 6E). From these data, we conclude that typical Rem2 signaling acts to regulate the number of dendritic segments both added and removed throughout early development. In addition, deletion of Rem2 results in a developmental trajectory dominated by changes in segment number as opposed to segment length (compared Fig 3E to 6E).

### Rem2 Bi-Directionally Regulates Segment Number in Distinct Developmental Epochs

After finding that Rem2 loss disrupts the developmental trajectory of basal arbors (Fig. 6B-E), we were interested to compare how the basal arbors of *Rem2^-/-^* cells differ from wildtype arbors at different ages. We found significant changes only during the early postnatal period (P7) and during the early visual critical period (P21), but not at other time points. Time points with no significant changes in any of the analyzed measures (P12, P16, and P30) have not been included in Figure 6. As suggested by our developmental results, we observe that *Rem2^-/-^* cells show changes in the number of dendritic segments (Fig. 6G), but not their length (Fig. 6F), in a developmentally-regulated fashion. Early in development (P7), we observed a significant decrease in the number of dendritic segments in *Rem2^-/-^* cells (Fig. 6G) that is absent at P12. In contrast, we observed an increase in the number of dendritic segments by our early critical period time point (P21, Fig. 6G). We were interested to see if this dysregulation of segment number resulted in a change in the total length of the arbor or represented instead a rearrangement of a set arbor length (Fig. 6H). While we observed a numerical increase in the mean arbor length of *Rem2^-/-^* cells at P21, this narrowly misses our criterion for statistical significance (p=0.051, α=0.05, Fig. 6H), suggesting that Rem2 primarily regulates the formation of segments as opposed to the amount of dendritic material composing the arbor. We also did not observe a significant change in the number of dendritic primary arbors at either P7 or P21 (Fig. 6I), suggesting that segments are respectively absent or added to existing primary arbors.

### Rem2 is an Experience-Dependent Negative Regulator of Segment Number

Rem2 function has previously been shown to be regulated by neuronal activity [51, 54, 67]. Therefore, we investigated whether visual experience was required for Rem2-dependent regulation of segment number. To test this, we dark-reared *Rem2 ^+/+^* and *Rem2^-/-^* mice from before eye opening until P21 and assessed segment number. We found that *Rem2^-/-^* cells reconstructed from dark-reared mice failed to increase segment number (Fig. 7C) compared to either *Rem2^+/+^* group (TR and DR) and had significantly fewer dendritic segments than *Rem2^-/-^* arbors from mice that had visual experience (Fig. 7C). These results indicate that one activity-dependent function of Rem2 is to oppose visual experience-dependent segment addition/branching during the visual critical period. Interestingly, *Rem2^-/-^* neurons lack an experience-dependent enhancement of segment length, and *Rem2^-/-^* neurons from both dark-reared and typically-reared animals exhibit lengths that resemble *Rem2 ^+/+^* neurons from typically-reared animals (Fig. 7B & D). These data indicate that Rem2 may normally act to oppose the growth of segment length in the absence of visual experience. Adding to the evidence that Rem2 mediates both segment number and length, *Rem2^-/-^* cells show a complementary set of dark rearing-associated changes to those observed in *Rem2^+/+^* cells at P21 (Fig. 7F) such that *Rem2^-/-^* cells show changes only in parameters of number whereas *Rem2^+/+^* cells show changes only in parameters of length. Interestingly, overall path length is similar in dark-reared and typically-reared mice, with neither *Rem2^+/+^* nor *Rem2^-/-^* cells showing a change regardless of rearing regime.

**Figure 7:**
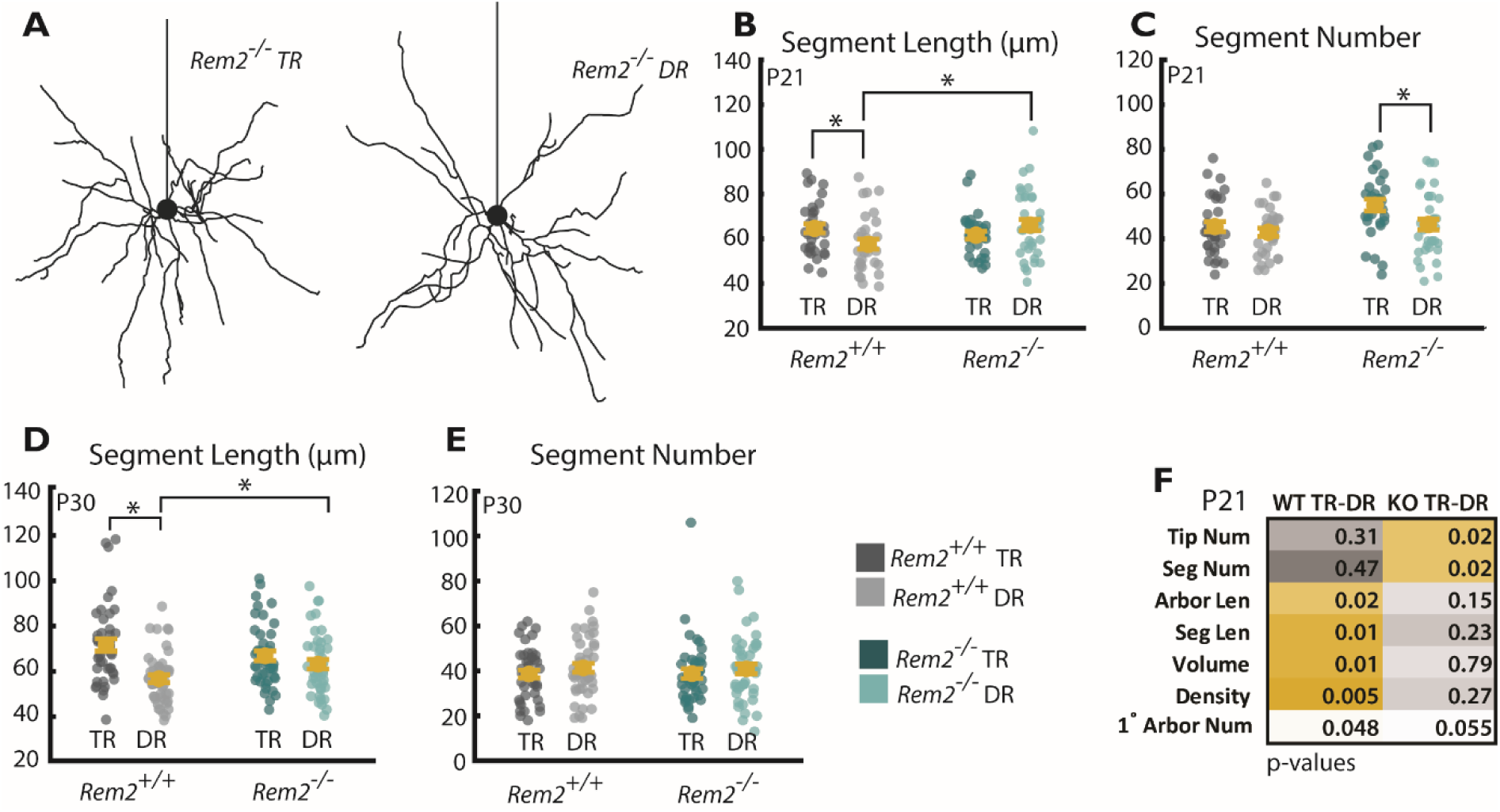
Rem2 regulates segment number in an experience-dependent manner. **A)** Example *Rem2^-/-^* neurons with (left) and without (right) visual experience. **B)** Average segment length of *Rem2 ^+/+^*and *Rem2^-/-^* neurons exposed either to typical rearing conditions or dark rearing from before eye opening and sampled at P21. **C)** Number of dendritic segments of *Rem2 ^+/+^* and *Rem2^-/-^* neurons exposed to either typical rearing conditions or dark rearing from before eye opening sampled at P21. **D)** Average segment length of *Rem2 ^+/+^* and *Rem2^-/-^* neurons exposed either to typical rearing conditions or dark rearing from before eye opening and sampled at P30. **E)** Number of dendritic segments of *Rem2 ^+/+^*and *Rem2^-/-^* neurons exposed to either typical rearing conditions or dark rearing from before eye opening sampled at P30. **F)** Summary of p-values from all comparisons when *Rem2 ^+/+^* and *Rem2^-/-^*are exposed to dark rearing from before eye opening until P21. Yellow boxes indicate significant results after multiple comparisons correction. Error bars are mean ± SEM. Yellow dots indicate means. Teal dots (*Rem2^-/-^*) and grey dots (*Rem2 ^+/+^*) indicate single neurons. For all *Rem2^-/-^*measurements: P7 TR N=21 cells, 2 mice; P12 TR N=47 cells, 4 mice; P16 TR N=44 cells, 4 mice; P21 TR N=30 cells, 3 mice; P30 TR N=43 cells, 4 mice; P21 DR N= 36 cells, 4 mice. For all *Rem2^+/+^* measurements: P7 TR N=24 cells, 2 mice; P21 TR N=33 cells, 3 mice; P21 DR N=32 cells, 4 mice (* indicates P<0.05, Wilcoxon rank-sum test).

### Acute, Sparse Deletion of Rem2 during the Visual Critical Period Alters Arbor Directionality

Our results indicate that Rem2 primarily contributes to arbor development and plasticity through establishing the number of segments in the dendritic arbor with modest impact on the length of segments or the total length of dendritic material composing the arbor. However, our *Rem2^-/-^* mice are constitutive null for *Rem2*, which eliminates our ability to manipulate Rem2 expression with any spatial or temporal control. Additionally, because of the constitutive nature of our *Rem2^-/-^* mouse model, we are unable to determine whether the observed changes are cell-autonomous or may originate from network-level changes in *Rem2^-/-^*cortices. To overcome these limitations, we crossed Rem2 conditional knockout mice (*Rem2^flx/flx^*) to Ai9.TdTomato (Tdt^flex/flex^) reporter mice that express the fluorophore TdTomato when Cre recombinase is present. We performed small, dilute injections of an AAV expressing GFP-tagged Cre recombinase into the visual cortex *of Rem2^+/+^;Tdt^flex/flex^* and *Rem2^flx/flx^;Tdt^flex/flex^* at P20 (Fig. 8A-C) enabling us to visualize sparse populations of neurons (Fig. 8A) and reconstruct whole arbors from just the TdTomato reporter fill (Fig. 8B). We sampled arbors approximately every two days starting at three days post injection (3dpi) and continuing until the peak of the critical period (11 dpi, P31; Fig. 8C). We were unable to reliably locate cells with sufficient TdTomato expression for reconstruction at 3 dpi, so this group has been excluded from further analyses.

**Figure 8:**
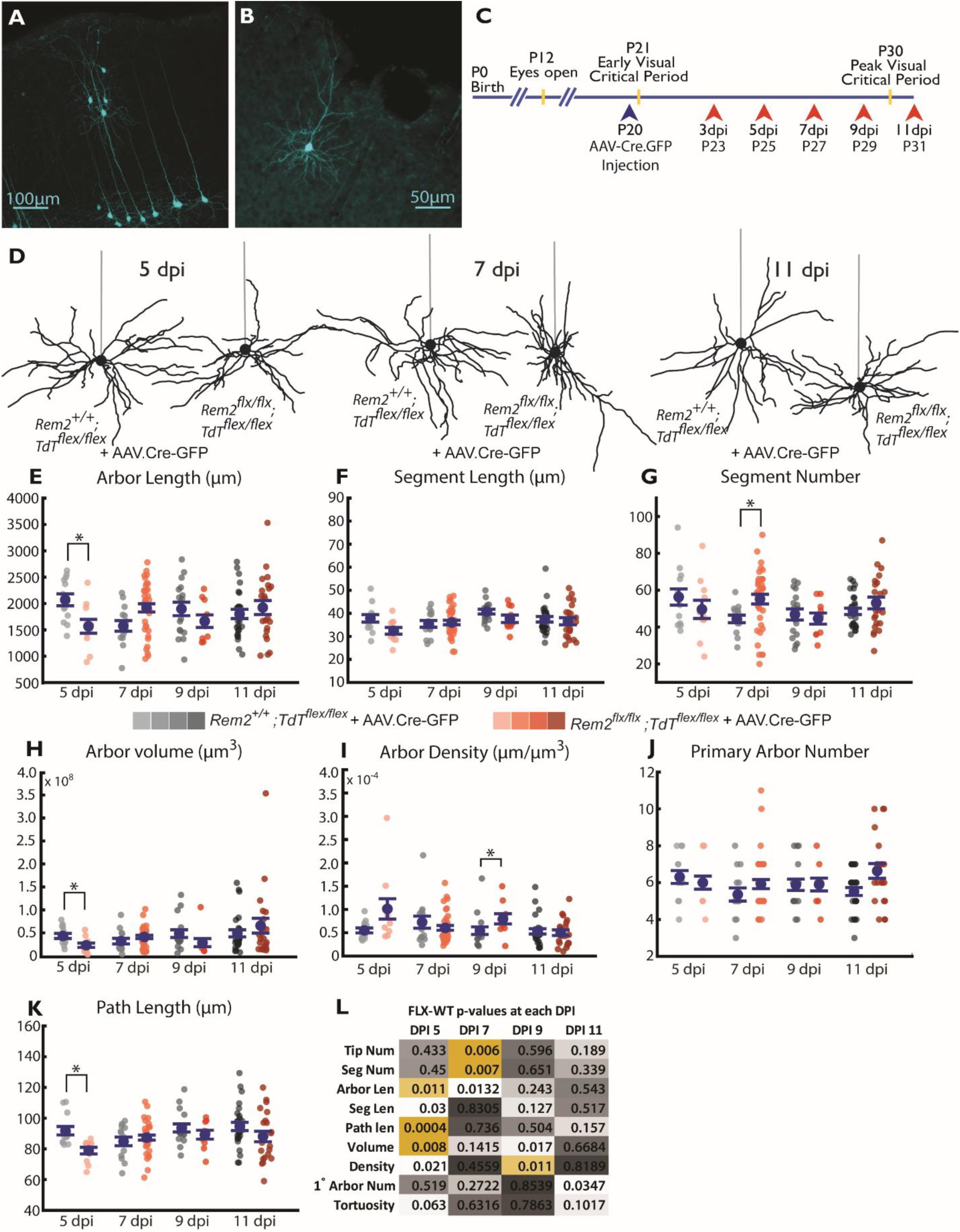
Acute Deletion of Rem2 results in many transient changes in basal arbor architecture. **A)** Example sparse population of wildtype neurons infected with AAV.Cre-GFP in primary visual cortex expressing Ai9.TdTomato reporter (pseudocolored cyan). **B)** Example single wildtype neuron expressing Ai9.TdTomato reporter (pseudocolored cyan). Entire dendritic arbor is visible and amenable to reconstruction. **C)** Timeline for AAV.Cre-GFP injection (blue arrow head) and morphological sample collection schedule (coral arrow heads). **D)** Example L2/3 pyramidal neurons in the primary visual cortex of *Rem2^+/+^;Tdt^flex/flex^* + AAV.Cre-GFP and *Rem2^flx/flx^;Tdt^flex/flex^* + AAV.Cre-GFP mice sampled 5 days post- injection (5dpi), 7 days post-injection (7dpi), or 11 days post-injection (11dpi). Straight gray lines diagram the unreconstructed apical dendrite to indicate the orientation of cells. **E)** Length of the complete basal arbor of *Rem2^+/+^;Tdt^flex/flex^* + AAV.Cre-GFP and *Rem2^flx/flx^;Tdt^flex/flex^* + AAV.Cre-GFP neurons sampled at 5dpi, 7dpi, 9dpi and 11dpi. **F)** Average segment length of neurons from *Rem2^+/+^;Tdt^flex/flex^*+ AAV.Cre-GFP and *Rem2^flx/flx^;Tdt^flex/flex^* + AAV.Cre-GFP mice sampled at 5dpi, 7dpi, 9dpi, and 11dpi. **G)** Segment number of neurons from *Rem2^+/+^;Tdt^flex/flex^* + AAV.Cre-GFP and *Rem2^flx/flx^;Tdt^flex/flex^* + AAV.Cre-GFP mice sampled at 5dpi, 7dpi, 9dpi, and 11 dpi. **H)** Volume of the complete basal arbor of *Rem2^+/+^;Tdt^flex/flex^* and *Rem2^flx/flx^;Tdt^flex/flex^* + AAV.Cre-GFP neurons sampled at 5 dpi, 7dpi, 9dpi and 11dpi. **I)** Density of the complete basal arbor of *Rem2^+/+^;Tdt^flex/flex^*+ AAV.Cre-GFP and *Rem2^flx/flx^;Tdt^flex/flex^* + AAV.Cre-GFP neurons sampled at 5 dpi, 7dpi, 9dpi and 11dpi. **J)** Number of basal dendritic primary arbors of *Rem2^+/+^;Tdt^flex/flex^*+ AAV.Cre-GFP and *Rem2^flx/flx^;Tdt^flex/flex^* + AAV.Cre-GFP neurons sampled at 5 dpi, 7dpi, 9dpi and 11dpi. **K)** Average path length of neurons from *Rem2^+/+^;Tdt^flex/flex^* + AAV.Cre-GFP and *Rem2^flx/flx^;Tdt^flex/flex^* + AAV.Cre-GFP mice sampled at 5dpi, 7dpi, 9dpi, and 11dpi. **L)** Summary of p-values from all comparisons between *Rem2^+/+^;Tdt^flex/flex^*+ AAV.Cre-GFP and *Rem2^flx/flx^;Tdt^flex/flex^* + AAV.Cre-GFP neurons at each of the sampled dpi time points with significant results following Bonferroni correction shown in yellow. Arbor characteristics impacted by Rem2 deletion are not conserved across ages/time points. Error bars are mean ± SEM. Blue dots indicate means and gray/coral dots indicate single neurons. For all *Rem2^flx/flx^;Tdt^flex/flex^* + AAV.Cre-GFP measurements: 05dpi N=11 cells, 4 mice; 07dpi N=38 cells, 7 mice; 09dpi N=10 cells, 4 mice; 11dpi N=22 cells, 4 mice. For all *Rem2^+/+^;Tdt^flex/flex^*+ AAV.Cre-GFP measurements: 05dpi N=13 cells, 5 mice; 07dpi N=14 cells, 5 mice; 09dpi N=17 cells, 4 mice; 11dpi N=27 cells, 6 mice. (* indicates P<0.05, Wilcoxon rank-sum test).

Qualitative observation of *Rem2^flx/flx^;Tdt^flex/flex^* + AAV.Cre-GFP arbors at all dpi reveals dendrites that are often extensively self-overlapping and have unusual distributions around the soma (Fig. 8D). Quantification of these arbors revealed complex, varied remodeling of arbors following acute Rem2 deletion (summarized in Fig. 8L) that occur as early as 5 days after Rem2 knockout (5 dpi) and are transient on a timescale of days. The total arbor length of *Rem2^flx/flx^;Tdt^flex/flex^* + AAV.Cre-GFP cells is significantly decreased compared to *Rem2^+/+^;Tdt^flex/flex^* + AAV.Cre-GFP cells at 5 dpi and increased at 7 dpi, with no significant change at 9 dpi or 11 dpi (Fig. 8E). In contrast to previous results with constitutive Rem2 knockout, we observed compromised age-dependent arbor stability that is present across both measures of segment length and number in the context of visual experience that are highly dependent on the duration of Rem2 loss or, perhaps, animal age (Fig. 8L). We found that segment number is increased compared to *Rem2^+/+^;Tdt^flex/flex^* + AAV.Cre-GFP cells at 7 dpi in *Rem2^flx/flx^;Tdt^flex/flex^* + AAV.Cre-GFP (Fig. 8G), but that only segment length is decreased at 5 dpi without a significant change in segment number (Fig. 8F). By 11 dpi, we found no significant changes in either arbor-centric or tissue-centric parameters, perhaps indicating that arbors have stabilized following Rem2 deletion. From these results, we conclude that acute Rem2 deletion in individual cells during the critical period leads to altered arbor stability arising from transient, bidirectional changes in several parameters of arbor complexity.

Upon our initial observations of *Rem2^flx/flx^;Tdt^flex/flex^* + AAV.Cre-GFP neurons, we noticed that several of the neurons exhibited highly irregular basal dendrite orientation and directionality (Fig. 9A). For example, some neurons had the majority of paths directed to either the medial or lateral side of the soma (see Fig. 9A top) instead of originating from the basal surface of the soma, as is most typical in wildtype arbors (Fig. 4). Thus, we became interested in comparing the degree to which *Rem2^flx/flx^;Tdt^flex/flex^* + AAV.Cre-GFP express directionality and whether the directionality of these cells is similar to wildtype neurons.

**Figure 9:**
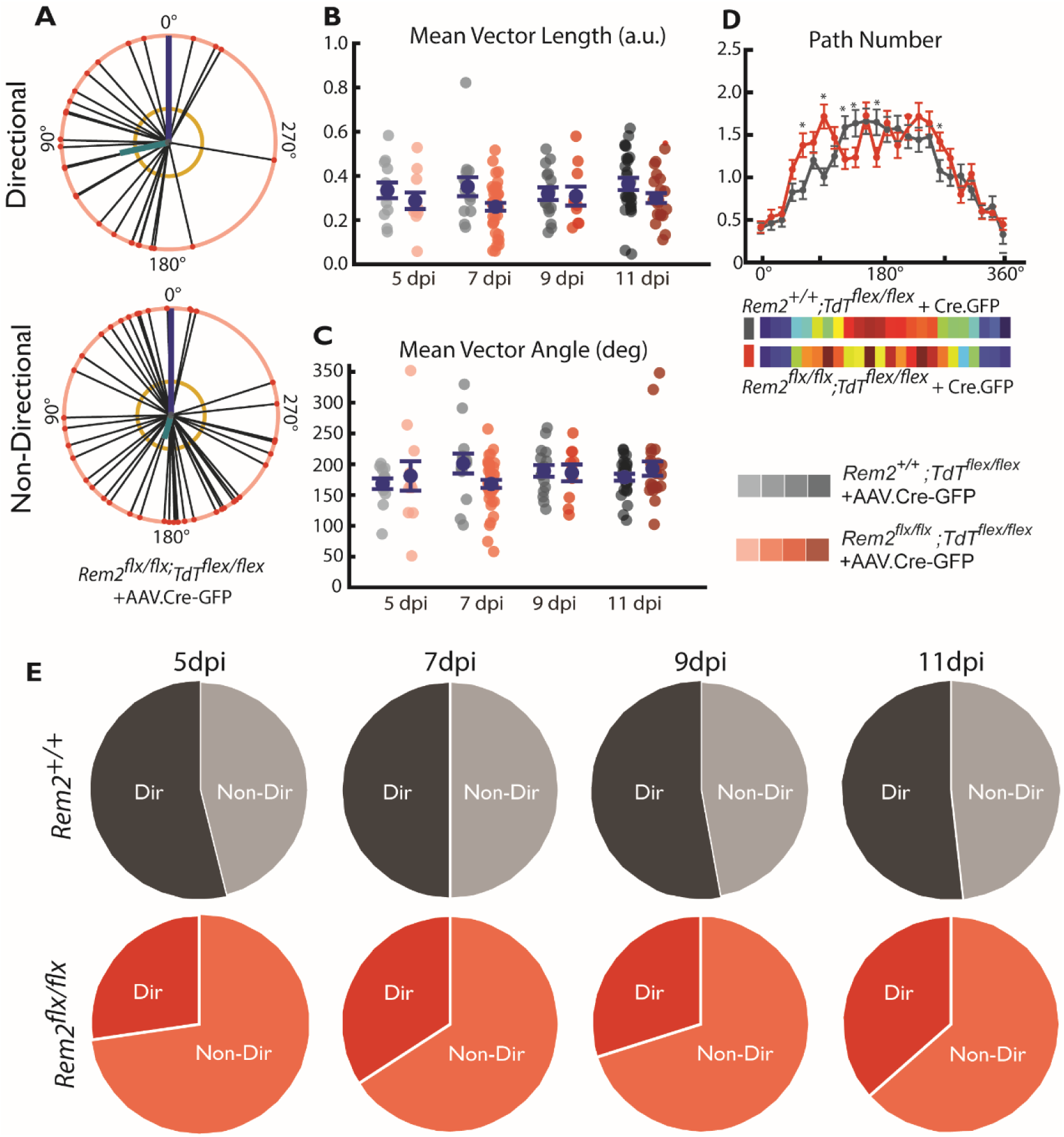
Rem2 promotes basal dendrite coronal polarity. **A)** Example directional (top) and non-directional (bottom) basal arbors sampled from *Rem2^flx/flx^;Tdt^flex/flex^* + AAV.Cre-GFP mice in the coronal viewpoint. Each arbor is schematized as soma (center dot) and soma-to-tip vectors (lines with dots at tip) arranged on a unit circle with apical dendrite diagramed as blue line. Thick teal line indicates the mean directional vector and yellow circle indicates length of mean directional vector required to declare the arbor directional by the Raleigh test. **B)** Length of the mean vector computed for *Rem2^+/+^;Tdt^flex/flex^* + AAV.Cre-GFP and *Rem2^flx/flx^;Tdt^flex/flex^* + AAV.Cre-GFP neurons sampled at 5dpi, 7dpi, 9dpi and 11dpi. **C)** Angle of the mean vector computed for *Rem2^+/+^;Tdt^flex/flex^* + AAV.Cre-GFP and *Rem2^flx/flx^;Tdt^flex/flex^* + AAV.Cre-GFP neurons sampled at 5dpi, 7dpi, 9dpi and 11dpi. D) Number of paths per radial bin (radial Sholl) for *Rem2^+/+^;Tdt^flex/flex^* + AAV.Cre-GFP and *Rem2^flx/flx^;Tdt^flex/flex^* + AAV.Cre-GFP neurons pooled across all ages (top) and heat map of the same data (bottom) * indicates P<0.05 by 1-Way ANOVA followed by Scheffe Test. **E)** Fraction of neurons meeting the Raleigh Test criterion for significant directionality of the basal arbor is noticeably decreased (∼20% decrease) in *Rem2^flx/flx^;Tdt^flex/flex^*+ AAV.Cre-GFP neurons at all sampled ages. Error bars are mean ± SEM. Blue dots indicate means and gray/coral dots indicate single neurons. For all *Rem2^flx/flx^;Tdt^flex/flex^*+ AAV.Cre-GFP measurements: 05dpi N=11 cells, 4 mice; 07dpi N=38 cells, 7 mice; 09dpi N=10 cells, 4 mice; 11dpi N=22 cells, 4 mice. For all *Rem2^+/+^;Tdt^flex/flex^*+ AAV.Cre-GFP measurements: 05dpi N=13 cells, 5 mice; 07dpi N=14 cells, 5 mice; 09dpi N=17 cells, 4 mice; 11dpi N=27 cells, 6 mice.

While we saw no significant difference in either the length of the mean vector (Fig.9B) or the angle of this vector (Fig.9C) when we considered coronal directionality we did observe a marked change in the number of neurons reaching the Raleigh criterion for significant directionality (Fig.9E). In *Rem2^+/+^;Tdt^flex/flex^* + AAV.Cre-GFP neurons, about 50% of the reconstructed cells achieved significant directionality consistently across the observed time points (see also Fig. 4). In contrast, only about 25-35% of *Rem2^flx/flx^;Tdt^flex/flex^* + AAV.Cre-GFP neurons meet this criterion, indicating that Rem2 would normally function to promote a more isotropic distribution of basal dendrites in the coronal plane. When we instead considered tangential directionality (Fig. S2), a different pattern of results emerges. We observe little significant Rem2-dependent regulation of tangential directionality and orientation. However, the 7 dpi time point shows that *Rem2^flx/flx^;Tdt^flex/flex^* + AAV.Cre-GFP neurons have decreased tangential directional vector length compared to their wildtype littermates (Fig. S2B). Taken together with our findings concerning coronal polarity, these results identify a new role for Rem2 in regulation of dendritic architecture and identify a cell-autonomous molecular regulator of this little-studied component of dendritic complexity [68].

These experiments reveal that Rem2 is required for the continued, active maintenance of the basal arbor during a time that previous results, including our own, would suggest arbors have reached stable maturity. Strikingly, notable changes occur in as little as 5 days after Cre injection and subsequent Rem2 deletion, highlighting the rapidity with which remodelling occurs in arbors that had previously developed normally. Following from this idea, our results challenge the notion that a dendritic arbor undergoing no observable net change is devoid of regulation– in contrast, it may instead be the case that an equilibrium of regulatory forces maintains the arbor and that removal of one regulator, such as Rem2, can disrupt the structure of an outwardly stable arbor.

## Discussion

### Summary

Dendrite morphogenesis is a complex developmental process subject to exquisitely precise molecular regulation. Neurons must find their proper location in the tissue [24, 25, 69, 85–87], polarize neurites into dendrites and axons [88, 89], and elaborate their dendritic arbor in adherence to the structure of the neural circuit of which they find themselves a part [87, 90–96]. Proper patterning of dendrites, for example, being the appropriate size and shape to tile functionally relevant spaces, is a key component to the development of neural circuits [87, 90–98]. Conversely, the dendritic arbor is shaped by ongoing circuit activity both in the format of early pre- and postnatal spontaneous activity as well as through sensory input in later postnatal development as we have shown here and numerous research groups have shown previously in other model systems. Many regulators of this process have been identified across a variety of species, though many of the details remain to be uncovered.

This study establishes a cohesive, detailed description of postnatal dendritic arbor development in mouse visual cortex with and without the influence of sensory experience. We establish that proper dendritic development is a carefully orchestrated process involving both experience-dependent and experience-independent components that establish distinct aspects of the mature arbor. Our results suggest that gross structure of basal dendritic arbors (number of primary neurites, number of segments, 85% of dendritic segment length) arises independently of sensory experience, but that sensory experience is required to achieve full dendritic segment length. Additionally, we characterize basal arbor directionality and orientation for the first time and show that basal dendrites tend to avoid the area surrounding the apical dendritic arbor.

Furthermore, we establish that Rem2 is an age- and experience-dependent regulator of this process, primarily impacting segment number as opposed to segment length. However, acute loss of Rem2 reveals a complex role for Rem2 in actively regulating many parameters of the basal arbor including both dendrite length and number parameters. Finally, we show for the first time that Rem2 promotes basal dendrite directionality with Rem2 deletion resulting in an approximately 20% decrease in significantly directional basal arbors.

### A Program of Dendritic Growth in the Visual Cortex

Prior to the results presented here, the works describing the maturation of basal arbors in rodent visual cortex were limited by the lack of digital 3-D reconstruction pipelines available at the time they were produced [36, 37]. These works were unable to differentiate between possible alternative growth programs of dendritic arbors, such as whether arbor growth proceeds through the addition of new segments combined with concomitant increases in segment length or whether length or segment additions alone dominate growth. We found that, with typical visual experience, arbor length increases primarily through the outgrowth of existing dendritic segments as opposed to the addition of new segments or more primary arbors. Consistent with previous works [37, 69], we found that the number of primary arbors is established very early in development, before postnatal day 7, and is stable throughout the time period we investigated. Likewise, the number of dendritic segments undergoes some modification throughout development, primarily a decrease in number near eye opening, but contributes relatively little to net arbor growth. In contrast, we found that increasing length of dendritic segments accounts for the vast majority of the increase in arbor length. These results suggest a separation in the regulation of segment length and number as demonstrated by separable patterns of developmental change as well as differential regulation by Rem2.

### Experience-Dependent and Experience-Independent Arbor Growth

Some of the earliest events in dendrite morphogenesis are determined by transcriptional programs, in which expression of a specific transcription factor influences numerous components of neuronal identity, including morphology of dendrites and axons [69, 94, 99–102]. For example, ectopic expression of Fezf2, a necessary transcription factor for L5 corticofugal projection neuron identity, in L2/3 and L4 neurons is sufficient to alter dendritic morphology, axonal targeting, and synapse localization *in vivo* [100, 101, 103]. However, not all characteristics of the arbor are determined in this way; refinement of the arbor also involves a dynamic process of growth and pruning, each with known molecular regulators. Work from the Cline lab among others have established numerous molecular regulators of dendritic outgrowth [46, 48, 49, 62–65, 70, 71, 104], including many pathways directly connected with circuit activity-dependent mechanisms promoting dendrite outgrowth [42, 46, 51, 66, 71, 105, 106]. The canonical model of this process, the Synaptotrophic Hypothesis, suggests that dendrites grow toward areas that have active inputs and are stabilized by the presence of maturing synaptic contacts [72].

Sensory experience plays a key role in the development of sensory cortices, as evidenced by decades of work describing experience-dependent maturation of receptive field properties and circuit function in the visual cortex of rodents, carnivores, and non-human primates. Interestingly, visual experience and maturation of visual circuits has already been connected to increases in dendritic complexity in some vertebrate systems, such as the *X.Laevis* tadpole optic tectum [46]. Our results suggest that a similar connection between visual experience and dendritic elaboration exists in mammalian sensory systems as well, but with a greatly diminished influence of visual experience on the size and shape of dendritic arbors. We found that visual experience promotes slightly increased arbor length and slightly increased segment length, with no significant impact on the number of dendritic segments. Visual experience also reshapes the spatial extent of arbors, leading to an increase in arbor volume and decrease in arbor density. However, our results demonstrate that the vast majority of arbor growth occurs independently from visual experience – at P21 and P30, only about an additional 15% of the total arbor length is added in the presence of experience (P21, WT DR arbor length is 83% of WT TR; P30 WT DR arbor length is 87% of WT TR). For example, we observe an increase in arbor length between P16 and P21, which is sustained at least through P30, even when mice have been dark reared since before eye opening. These results demonstrate separable experience-dependent and experience independent programs for arbor growth.

### Rem2 and Experience-Dependent Arbor Development

Our results here indicate that Rem2 may interact with several aspects of dendritic morphogenesis, including regulation of the shape of the basal arbor, the formation of new segments, the length of segments, as well as the territory the arbor occupies in the tissue. Rem2 was previously shown to negatively regulate dendritic complexity in cultured neurons [54, 55] and in *X.Laevis* optic tectum in an activity-dependent manner [51]. Here, we demonstrate both experience-dependent and experience-independent roles for Rem2 in regulating the basal arbor in the brain of mice during key developmental time points for visual cortex development. Our results demonstrate that postnatal arbor development of *Rem2^-/^*^-^ neurons is largely dominated by changes in the number of dendritic segments in contrast to changes in the length of dendritic segments, which accounts for the majority of arbor growth in wildtype cells. Further, and in contrast to wildtype arbors, *Rem2^-/-^* arbors demonstrated no change in segment length but increased segment number when visual experience was present. These results reveal two key roles of Rem2 in regulating the basal arbor. First, we can conclude that Rem2 acts to promote segment length growth in the presence of visual experience – when Rem2 was present, we saw increased segment length in neurons sampled from typically-reared mice compared to dark-reared mice; however, this increase is absent in *Rem2^-/-^* arbors with both typically-reared and dark-reared segments having similar length to those of typically-reared wildtype mice. Second, Rem2 functions to restrict the number of dendritic segments in response to visual experience. These results suggest that visual experience promotes a complex growth program for basal dendritic arbor maturation, but that specific negative regulators, such as Rem2, sculpt the developmental process into a length-dominated growth program.

### Rem2 and Maintenance of Established Dendritic Arbors

Here, we show that acute loss of Rem2 during the critical period leads to numerous changes in dendritic arbor structure that cannot easily be classified into one (or even a few) categories of changes. Perhaps most strikingly, these changes arise rapidly within only a few days after Rem2 deletion and impact arbors that had previously developed normally in intact cortical circuits. Furthermore, these findings strongly suggest ongoing, finely balanced regulation of dendritic architecture, even at an age (P21-P30) when dendritic arbors are observed to be relatively stable in our wildtype dataset (Fig. 4). For this reason, we conclude that Rem2 may regulate the ability of the arbor to maintain its structure in the face of changes in network activity and other intrinsic regulators of dendritic complexity. In vertebrate systems [42-44, 46-49, 62, 63, 65, 70-72] and in other regions of the mammalian brain [73, 74], changes in sensory experience or behavioral state, such as experiences a stressful situation, have been shown to alter dendritic complexity. It follows that a negative regulator of this process, a stabilizing molecule, would be useful for restricting these changes and acting as a “brake” on runaway remodeling.

It is interesting to consider whether dendritic stability in Rem2 conditional knockout neurons arises via direct interactions with factors that alter the dendritic arbor or rather arises as a secondary consequence of a Rem2-mediated loss of synaptic stability. It has long been hypothesized that maturation of synaptic contacts onto the arbor stabilizes dendritic segments and spares these segments from pruning [72, 75], such that increases in synapse number should lead to growth of the dendritic arbor. While much experimentation remains to be done in order to explore a possible connection between synaptic and dendritic stability in the context of Rem2 function, some of our recent work may provide a hint. When Rem2 is deleted during the critical period, the same time period in which we observe dendritic instability here, changes in miniature excitatory post-synaptic current (mEPSC) frequency and amplitude were unchanged 4 days after Rem2 deletion, but by 10 days post-deletion a 25% decrease in mEPSC frequency was observed [76]. Here, we observed arbor changes as early as 5 days post-infection, suggesting that arbor changes may lead synaptic changes or occur simultaneously. Future work using live imaging of dendrite and synapse dynamics in the context of Rem2 deletion could help to rigorously establish whether arbor changes occur earlier than, later than, or simultaneously with synaptic changes.

### The Many Functions of Rem2

Previous work has established that Rem2 plays a multi-faceted role in regulating cortical development, including functioning as a positive regulator of synapse formation [51, 55, 76–78], a negative regulator of intrinsic excitability [76], and as a required molecule for complete expression of juvenile ocular dominance plasticity in the mouse [76]. Here, we have added to these findings evidence that Rem2 is an age- and experience-dependent regulator of dendritic complexity and promotes basal dendrite directionality. Taken together, this body of work constructs a complex network of Rem2-dependent processes. However, a major outstanding question remains – by what mechanism(s) does Rem2 regulate these many developmental processes?

One possible explanation comes from recent data showing that Rem2 is a potent inhibitor of Ca^2+^/calmodulin-dependent protein kinase II (CaMKII) [79]. CaMKII is perhaps one of the most intensely studied proteins in excitable cells, such as neurons and cardiomyocytes, and is a key regulator of Ca^2+^ signaling and homeostasis in these cells. In the brain, CaMKII has been implicated in myriad signaling cascades and is widely considered to be the key molecular regulator of memory formation as well as playing important roles in synapse formation, synaptic plasticity, and neuronal development. In the visual cortex of mice, CaMKII is involved in synapse pruning following visual deprivation [80]. Furthermore, CaMKII mutant mice also show decreased potential for ocular dominance plasticity [81] and Hebbian forms of synaptic plasticity [82]. Interestingly, CaMKII has also been implicated in the maturation of dendritic arbors [44, 47, 54, 83, 84]. Some reports demonstrate additional complexity of this regulation, indicating isoform-specific roles for CaMKII, with CaMKIIβ restricting dendritic complexity in cultured granule neurons independently of CaMKIIα [84].

From these many known functions of CaMKII, we see that direct, potent inhibition of this molecule may be an important part of maintaining cellular homeostasis at a variety of levels. Likewise, loss of such inhibition leading to increased CaMKII catalytic activity, such as by Rem2 deletion, could lead to widespread changes in cellular structure and function. This is consistent with the many alterations we observe in our past and current studies of Rem2 deletion [51, 54, 55, 67, 76–79]. Compellingly, the ability for Rem2 to inhibit CaMKII catalytic activity has already been linked to regulation of dendritic complexity [79]. We propose a model in which direct inhibition of CaMKII by Rem2 promotes morphological stability, the loss of which results in cascading instability of the dendritic arbor.

### Implications for models

These results offer a number of useful constraints for those building models of cortical development and the influence of experience. Basal arbor tortuosity hardly varies across animals or development and takes values close to 1 (median: 1.23, interquartile range: 0.09), so a “ball and stick” model of basal dendritic architecture is an excellent approximation. The number of primary neurites is determined very early (by P7) and is not dependent on visual experience, and the approximate number of dendritic branches is also determined early. Basal arbors do grow with age and experience (42% increase in arbor length from P16 to P30) but grow only a little bit more with visual experience (15%) than without visual experience. These observations indicate that models that leave out an experience-dependent component of increases in basal dendritic length would not be missing too much, and models could instead focus on experience-dependent contributions of active channels and dynamic synapses over relatively fixed dendritic lengths.

## Conclusion

Here, we report that dendritic development in the mouse visual cortex proceeds through both experience-dependent and experience-independent mechanisms. In contrast to previous reports from the *X.Laevis* visual system, we find that experience plays a nuanced role in arbor refinement, and largely drives arbor growth through small increases in segment length rather than in increasing the number of segments. We also show that Rem2 is a key regulator of this process with both experience-dependent and experience-independent roles. Furthermore, we show that Rem2 is a cell-autonomous regulator of basal arbor stability and directionality, a new function for this molecule. We hypothesize that these functions of Rem2 depend on a recently-described [79] direct inhibition of CaMKII by Rem2, the loss of which leads to numerous downstream effects dependent on both network activity and age.

## Supplemental Figures and Captions

**Figure S1:**
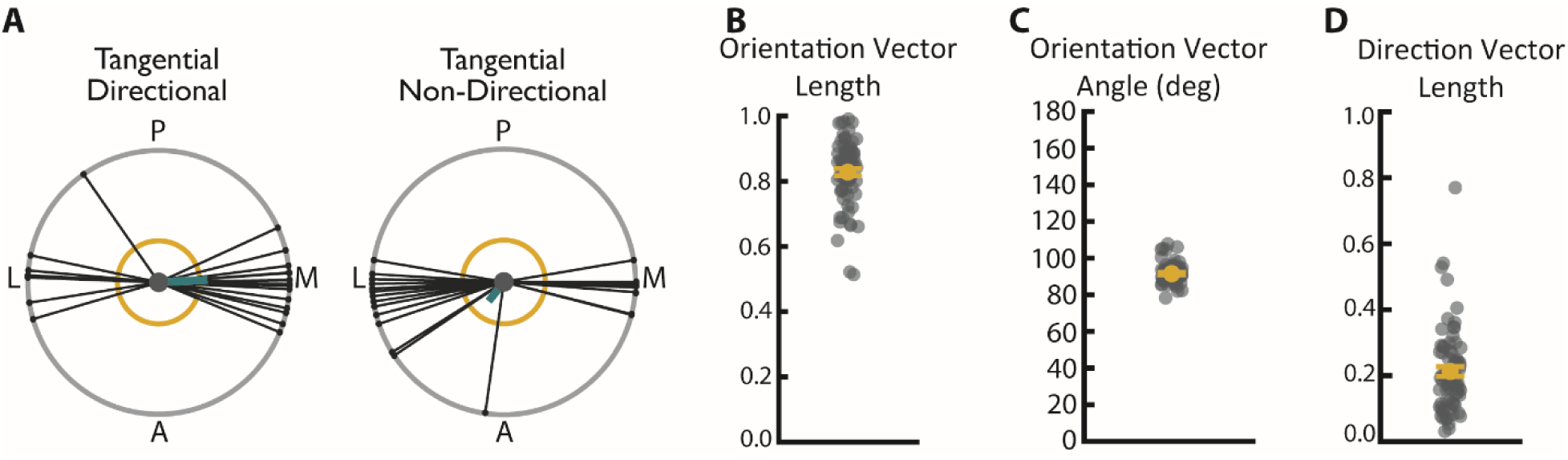
Wildtype arbors have oriented arbors in the tangential plane. **K)** Example directional (left) and non-directional (right) basal arbors sampled from wildtype mice in the tangential viewpoint. Each arbor is schematized as soma (center dot) and soma-to-tip vectors (lines with dots at tip) arranged on a unit circle. Thick teal line indicates the mean direction vector and yellow circle indicates length of mean direction vector required to declare the arbor directional by the Raleigh test. Posterior (P), anterior (A), medial (M) and lateral (L) are indicated around the diagrams. **L)** Length of the tangential normalized directional vector calculated from all basal paths. Larger values indicate greater directionality. **M)** Angle of tangential directional vector calculated from all basal paths. **N)** Length of the tangential normalized orientation vector calculated from all basal paths. Larger values indicated greater orientation.

**Figure S2:**
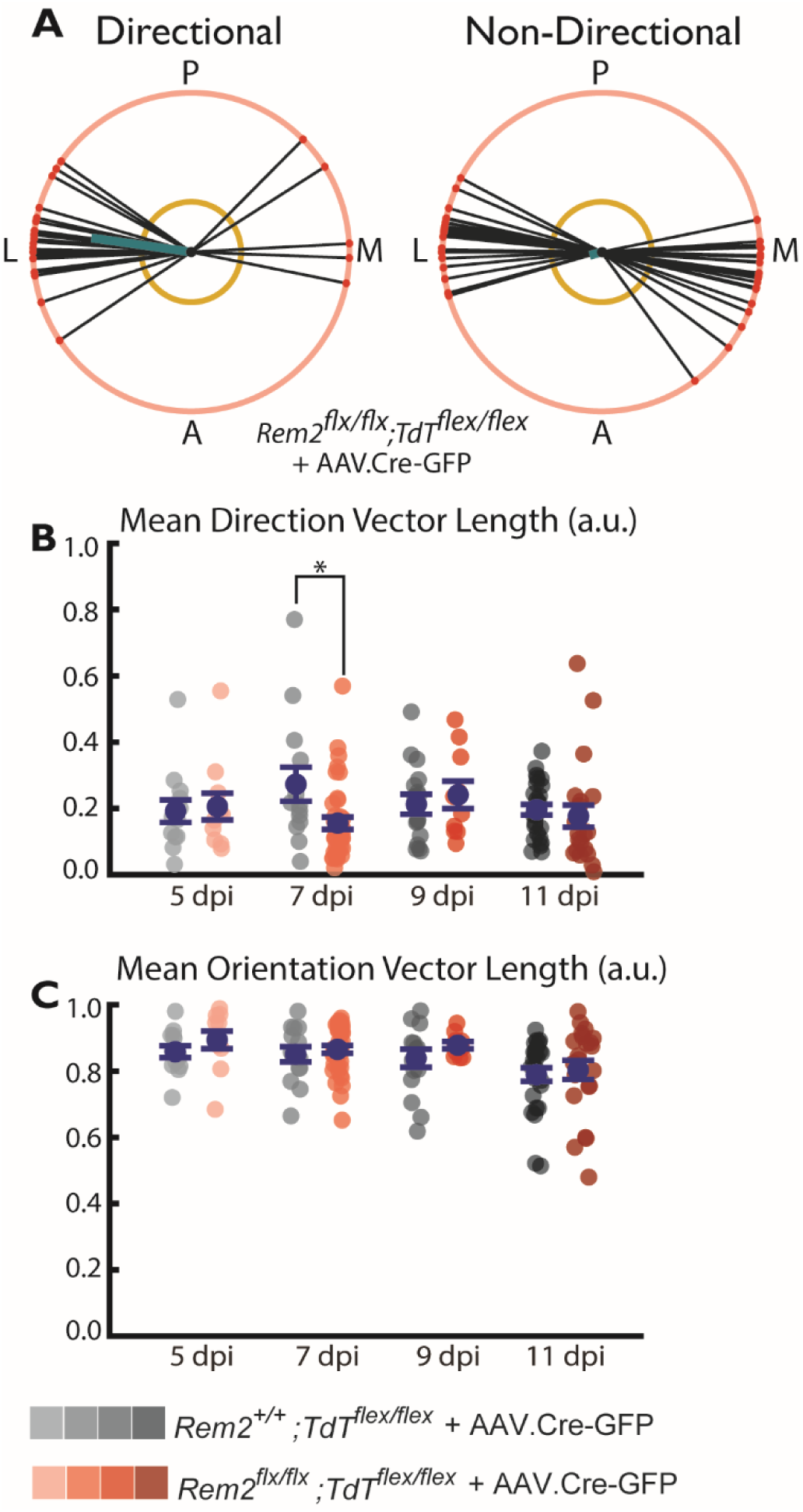
Rem2 promotes tangential directionality. **A)** Example polar (left) and non-polar (right) basal arbors sampled *Rem2^flx/flx^;Tdt^flex/flex^*+ AAV.Cre-GFP mice in the tangential viewpoint. Each arbor is schematized as soma (center dot) and soma-to-tip vectors (lines with dots at tip) arranged on a unit circle. Thick teal line indicates the mean direction vector and yellow circle indicates length of mean direction vector required to declare the arbor directional by the Raleigh test. Posterior (P), anterior (A), medial (M) and lateral (L) are indicated around the diagrams. **B)** Length of the mean direction vector computed for *Rem2^+/+^;Tdt^flex/flex^* + AAV.Cre-GFP and *Rem2^flx/flx^;Tdt^flex/flex^* + AAV.Cre-GFP neurons sampled at 5dpi, 7dpi, 9dpi and 11dpi. **C)** Length of the mean orientation vector computed for *Rem2^+/+^;Tdt^flex/flex^*+ AAV.Cre-GFP and *Rem2^flx/flx^;Tdt^flex/flex^* + AAV.Cre-GFP neurons sampled at 5dpi, 7dpi, 9dpi and 11dpi. Error bars are mean ± SEM. Blue dots indicate means and gray/coral dots indicate single neurons. For all *Rem2^flx/flx^;Tdt^flex/flex^* + AAV.Cre-GFP measurements: 05dpi N=11 cells, 4 mice; 07dpi N=38 cells, 7 mice; 09dpi N=10 cells, 4 mice; 11dpi N=22 cells, 4 mice. For all *Rem2^+/+^;Tdt^flex/flex^*+ AAV.Cre-GFP measurements: 05dpi N=13 cells, 5 mice; 07dpi N=14 cells, 5 mice; 09dpi N=17 cells, 4 mice; 11dpi N=27 cells, 6 mice. (* indicates P<0.05, Wilcoxon rank-sum test).

